# Excessive Ca^2+^-dependent ER-mitochondrial contact stabilization by EFHD1 drives liver injury

**DOI:** 10.64898/2026.02.13.705622

**Authors:** David R. Eberhardt, Emma C. Rekate, Yasmin B. Masini, Hannah E. Duron, David Mollinedo, Adrian M. Velarde, Devorah Stucki, Tara Price, Sandra H. J. Lee, Enrique Balderas, Neeraj K. Rai, Ashley R. Bratt, Anthony M. Balynas, Chris J. Stubben, Ryan Bia, Sudipa Maity, Nicolas Hartel, Xue Yin, Andrea Corbin, Anshu Kumari, Dung M. Nguyen, Daisuke Shimura, Vu D. Nguyen, Vishaka Vinod, Kamrul H. Chowdhury, Francisco Verdeguer, Joel Zvick, Patrice N. Mimche, Sihem Boudina, Stavros G. Drakos, Ademuyiwa S. Aromolaran, Sarah Franklin, Vivek Garg, Robin M. Shaw, William L. Holland, Scott A. Summers, Marcus G. Pezzolesi, Jared Rutter, Kimberley J. Evason, Dipayan Chaudhuri

**Author notes:** Corresponding author: Dipayan Chaudhuri, MD, PhD, Nora Eccles Harrison Cardiovascular Research and Training Institute University of Utah 95 S 2000 E Salt Lake City, UT 84112.

## Abstract

Metabolic-associated steatohepatitis (MASH) involves hepatocyte damage that cannot be explained solely by lipid accumulation. Here, to discover injury-specific pathways, we focused on a gene of uncertain function, *EF-Hand Domain Family Member D1* (EFHD1), identified in human genome-wide association studies of liver injury but not liver fat. We show that EFHD1, a Ca^2+^-dependent actin crosslinker, stabilizes endoplasmic reticulum–mitochondria contact sites (ERMCS), detecting spatiotemporal coincidence of inter-organellar proximity and ER Ca^2+^ release. During MASH, EFHD1 upregulation drives pathological mitochondrial fragmentation via excessive contact persistence. This structural failure promotes mitochondrial double-stranded RNA escape and activation of a maladaptive antiviral PKR-dependent stress response, a causal relationship also supported by Mendelian randomization in humans. Consequently, inhibiting EFHD1 in human and mouse models blunts hepatocyte damage. These findings identify EFHD1 as a Ca^2+^-dependent ERMCS stabilizer, reveal a hepatocyte-intrinsic injury pathway, and suggest EFHD1 inhibition as a therapeutic strategy.

## INTRODUCTION

The growing prevalence of metabolic dysfunction-associated steatotic liver disease and steatohepatitis (referred to hereafter as MASH) is an area of profound unmet clinical need(1). One approach for therapeutic and mechanistic insight has been to seek genes altering MASH risk(2, 3). Multiple genome-wide association studies (GWAS) for MASH have assessed either liver fat content or susceptibility to liver injury. Both types of analyses have repeatedly identified genes critical for hepatic lipid metabolism. However, lipid accumulation alone does not predict progression to MASH, and therapies inhibiting lipid metabolism have been hampered by difficulties (4–6). Therefore, to identify pathways that contribute to liver injury independently of lipid metabolism, we focus here on a gene of unknown function repeatedly identified in GWAS of liver enzymes, but not in GWAS measuring liver fat, *EF-Hand Domain Family Member D1* (EFHD1)(7–10).

EFHD1 is a 27-kDa nuclear-encoded protein that localizes to mitochondria and is expressed widely. Variation in the *EFHD1* locus has been consistently associated with serum levels of liver enzymes, markers of liver injury, across multiple populations(7–10). The variant most strongly associated with elevated serum liver enzymes increases liver-specific expression of *EFHD1* by encoding a better consensus sequence for hepatic transcription factors, including FOXA2, FOXO1, and HNF4A(11–13). However, modulating EFHD1 does not produce a consistent effect on glycolysis or fatty acid metabolism across different cell types(12–15). Thus, whether EFHD1 truly contributes to MASH progression, and by what mechanism, remain critical unanswered questions.

Here, using mouse and human models, we show that EFHD1 contributes to a central pathway for hepatocyte injury. Notably, this occurs without altering organismal energy balance or hepatic steatosis, consistent with human GWAS linking *EFHD1* to liver enzymes but not fat, and suggesting that the EFHD1-dependent injury is independent of the initial insult. Mechanistically, we identify that EFHD1, a Ca²⁺-dependent actin crosslinker, functions as a spatiotemporal coincidence detector, converting transient ER Ca²⁺ signals into mechanically persistent ER–mitochondria coupling. During metabolic stress, EFHD1 upregulation leads to pathological contact persistence and aberrant mitochondrial fission. These events promote cytoplasmic release of mitochondrial double-stranded RNA (mt-dsRNA) and activation of a maladaptive integrated stress response, a pathway normally reserved for antiviral defense. Finally, EFHD1 inhibition blunted hepatocyte injury, inflammation, and fibrosis across multiple metabolic and chemical liver disease models, suggesting it is a tractable target for selectively uncoupling liver injury from lipid accumulation.

## RESULTS

### EFHD1 is expressed in hepatocytes and increases during MASH

In humans, EFHD1 is preferentially expressed in hepatocytes, with enrichment in periportal hepatocyte clusters with high complement and immune pathway activation (**Fig. 1A-B, S1A**)(13, 16–20). Moreover, we found increased liver EFHD1 in humans with MASH (**Fig. 1C-D**). To investigate EFHD1 function further, we treated mice with three separate diets, a normal chow diet (10% fat), a high fat diet (HFD, 60% fat, 18% protein, 22% carbohydrate), which produces obesity but only mild liver injury, and a Gubra Amylin MASH diet (40% fat, 2% cholesterol, 20% fructose), which mimics hallmarks of human MASH injury including hepatocyte ballooning, inflammation, and fibrosis (**Fig. S1B**)(21, 22). For HFD and MASH, mice were started on the diet at 8 weeks of age for up to 28 weeks duration (36 weeks of age). EFHD1 increased 2-4-fold in wild-type (WT) animals on both HFD and MASH diets (**Fig. 1E-F, S1C-D**). Taken together, human and mouse studies suggest EFHD1 is expressed in hepatocytes and is upregulated during overnutrition.

**Figure 1.**
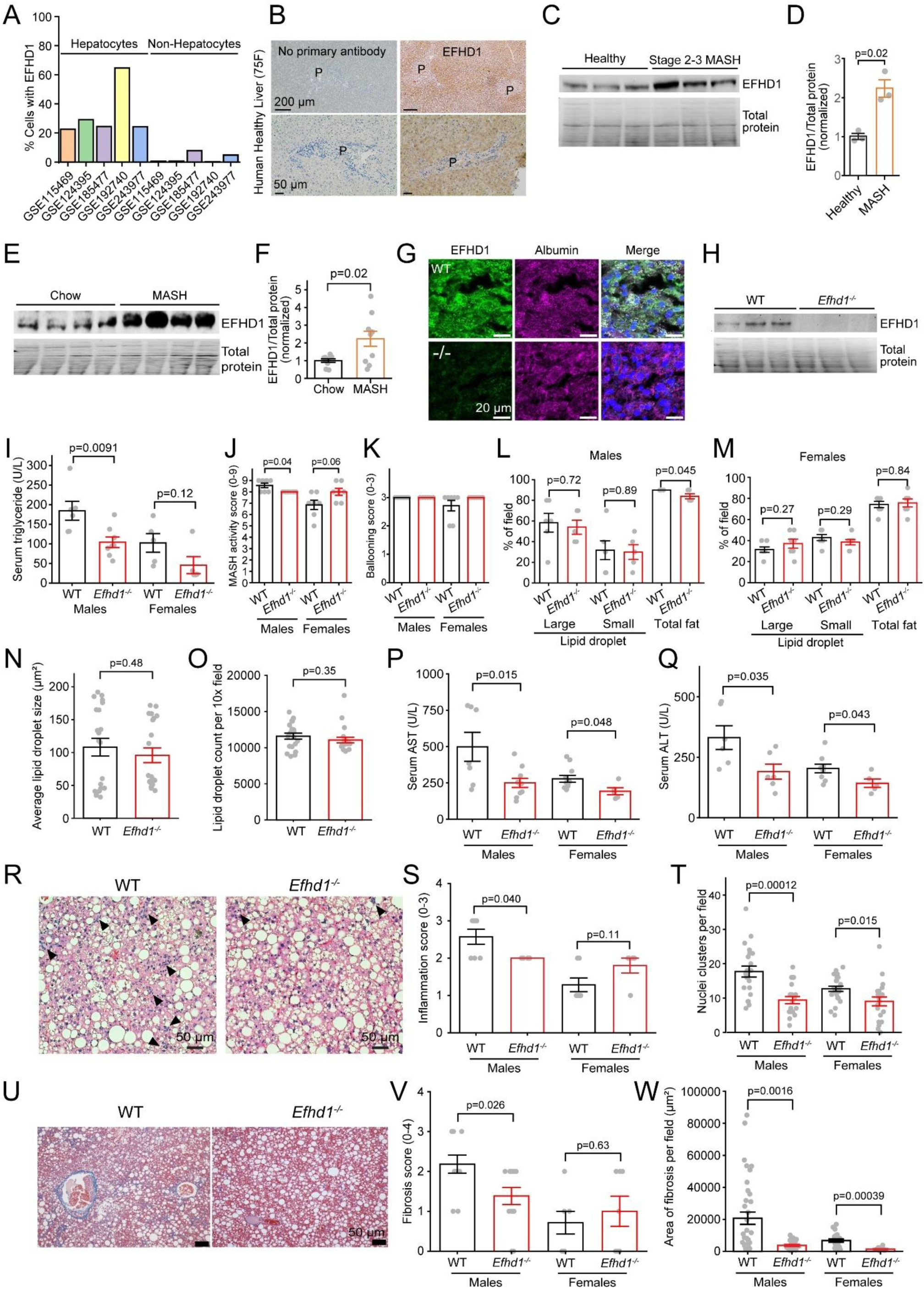
Loss of EFHD1 prevents hepatocyte injury in MASH. **A.** Summary of human liver single-cell RNA-seq studies. **B.** Immunohistochemistry showing EFHD1 is enriched around portal tracts (P) in healthy human liver (n=8). **C, D.** Western blot (C) and summary band quantification (D), human MASH (n=3). **E, F.** Western blot and summary, MASH diet mice (n=8). **G.** EFHD1 (green) co-expression with the hepatocyte marker, albumin (magenta) in WT or *Efhd1^-/-^* livers. In merge, nuclei are DAPI-stained. **H.** EFHD1 is absent in *Efhd1^-/-^*livers (n=3). **I-W.** Assays performed on mice following 28 weeks of MASH diet. **I.** Serum triglyceride levels. WT: males n=6, females n=5; *Efhd1*^-/-^: males n=8, females n=5. **J.** MASH activity score. **K.** Hepatocyte ballooning score. **L, M.** Histopathological assessment of liver fat. **N, O.** Direct measurement of lipid droplet size (N) and count (O). **P, Q.** Serum AST (P) and ALT (Q) measurements. **R.** Liver hematoxylin and eosin staining. Arrows, leukocyte clusters. **S.** Histopathological inflammation score. **T.** Liver leukocyte cluster counts. **U.** Liver Masson’s trichrome staining. Blue, fibrosis. **V.** Histopathological fibrosis score. **W.** Fibrotic area from liver micrographs. All clinical histopathological scoring in J, K, L, M, S, and V was assessed by liver pathologist (K.J.E.). For J-Q, S, V, WT: males n=7, females n=7; *Efhd1^-/-^*: males n=6, females n=7. For N, O, WT: n=21; *Efhd1^-/-^*: n=18. For T, W, WT: males n=21, females n=21; *Efhd1^-/-^*: males n=21, females n=21.Bars: mean ± SEM.

### Deletion of EFHD1 is protective in MASH without affecting energy balance

To investigate whether EFHD1 contributes to liver injury, we examined whole-body EFHD1 knockout mice (*Efhd1^-/-^*) (**Fig. 1G-H**)(23). The mice exhibited normal weight gain, body composition, feeding, activity, insulin resistance, and liver:body weight (**Fig. S1E-R**), establishing that loss of EFHD1 is well-tolerated and changes in hepatic physiology are independent of global energy balance.

We measured hepatic function after a 4-hour fast, assaying hepatic lipid metabolism by serum triglycerides, histopathological scoring by a pathologist blinded to genotype, and direct lipid droplet measurement (**Fig. 1I-O, S2A-N**)(24, 25). Notably, though *Efhd1^-/-^*mice had reduced hepatic lipid on a normal diet, these differences disappeared in the obesogenic diets, suggesting EFHD1 affects hepatic lipid metabolism indirectly and concordant with the absence of *EFHD1* in human liver fat GWAS.

In contrast, injury markers were markedly reduced in *Efhd1^-/-^* across diets, assessed both via serum liver enzymes and histology (**Fig. 1P-W, S2O-W**). Serum aspartate aminotransferase (AST) and alanine aminotransferase (ALT) levels in *Efhd1^-/-^* mice were about half of wild-type (**Fig. 1P-Q, S2O-R**). Liver inflammation and fibrosis were also reduced in males on both histopathological scoring and direct measurement (**Fig. 1R-T, S2S-W**). In *Efhd1^-/-^* livers, we also noted fewer inflammatory cells per cluster (**Fig. 1R, S2S**). Female mice are less injured by overnutrition(26), so they had minimal inflammation, though inflammatory clusters and fibrotic area were still reduced in MASH *Efhd1^-/-^* females (**Fig. 1T, W**). Fibrosis was minimal for chow and HFD and not scored (**Fig. S2T**). To summarize, loss of EFHD1 is protective during MASH progression due to reduced liver injury.

### EFHD1 is necessary for Ca^2+^-induced mitochondrial remodeling

Because EFHD1 localizes to the outer mitochondrial membrane (OMM), including ER-mitochondrial contact sites (ERMCS) (**Fig. S3A-B**), we tested its contribution to mitochondrial morphology(23, 27–30). We examined mouse hepatocytes within 24 hours of isolation, prior to de-differentiation (**Fig. S3C-E**), using MitoTracker Orange. Intriguingly, whereas WT mitochondria appeared bean-shaped, *Efhd1^-/-^*mitochondria were longer and spaghetti-shaped (**Fig. 2A-B**). MitoTracker Orange only labels polarized mitochondria, confirming this was not due to differences between injured versus healthy mitochondria. This difference was notable even in hepatocytes isolated from mice subject to MASH diet, which causes extreme mitochondrial fission (**Fig. 2C**). Moreover, the difference in mitochondrial length was also evident on transmission electron micrographs (TEM) (**Fig. 2D-F, S3F**). Finally, this effect was not hepatocyte-specific, and could be replicated in cultured *EFHD1^-/-^* HepG2 and HAP-1 cells, and could be rescued by re-expression of EFHD1 (**Fig. 2G-I, S3G-J**)(31).

**Figure 2.**
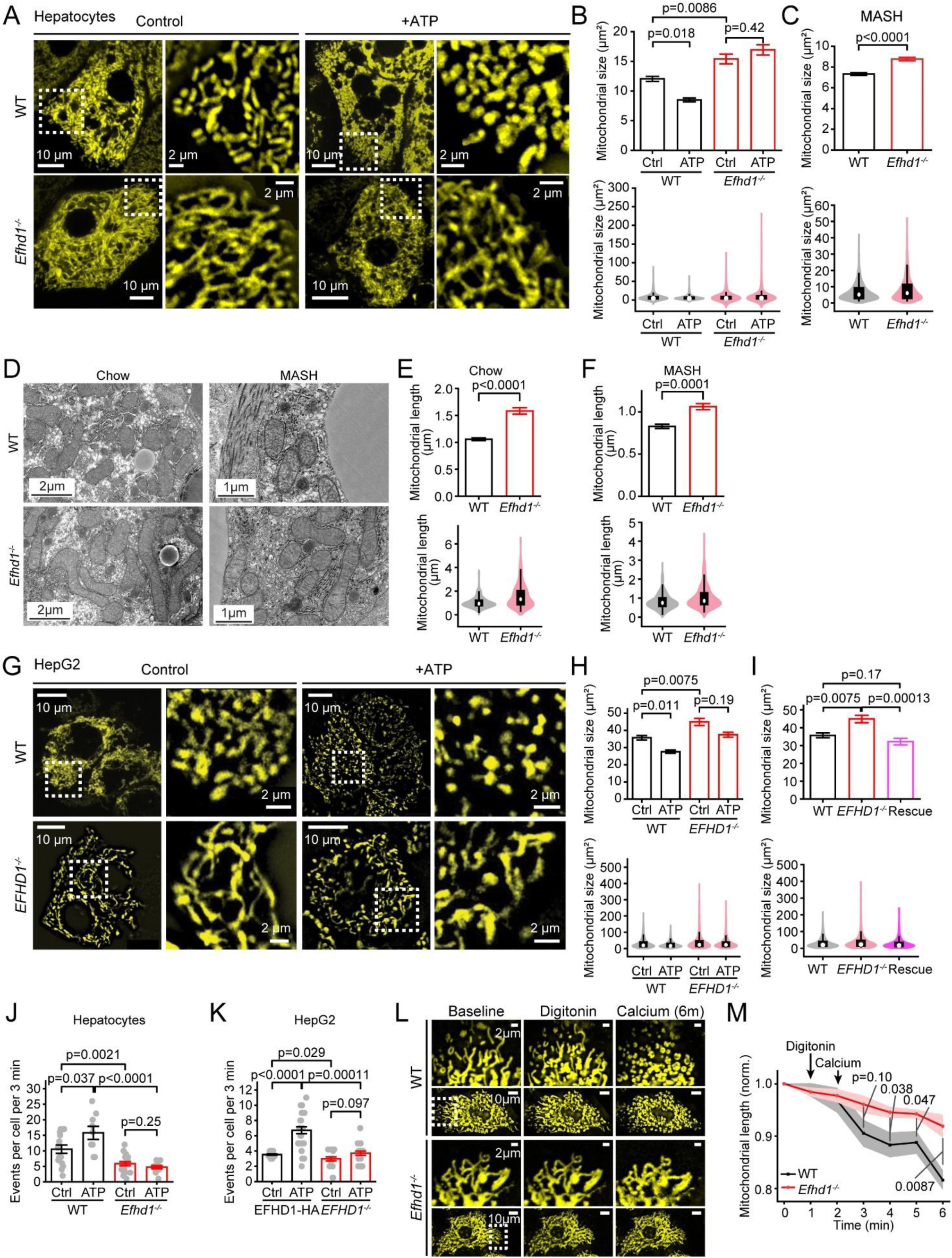
EFHD1 is necessary for Ca^2+^-induced mitochondrial remodeling. **A.** Representative hepatocytes, 200 nM MitoTracker Orange staining before (Control) or 5 minutes after treatment with 1 mM ATP (+ATP). Right, boxed insets at higher magnification. **B.** Automated iLastik analysis, hepatocyte mitochondrial size. Data is displayed as a bar chart (top) to show mean effects and as a violin plot (bottom) to show the full distribution. WT Ctrl n=10008 mitochondria from N=6 mice; WT ATP N=5, n=7740; *Efhd1*^-/-^ Ctrl N=7, n=6577; *Efhd1*^-/-^ ATP N=5, n=4160. **C**. Hepatocyte mitochondrial size after 28 weeks MASH diet. WT n=7071 mitochondria from N=3 mice, *Efhd1*^-/-^ Ctrl: N=3, n=4642. **D.** Representative TEM of mouse livers. **E, F.** Hepatocyte mitochondrial length from TEM. Chow: WT n=347 mitochondria from N=3 mice; *Efhd1^-/-^* N=3, n=273. MASH: WT N=3, n=274; *Efhd1^-/-^* N=3, n=294. **G.** As in (A), but for WT or *EFHD1^-/-^* HepG2 cells. **H.** As in (B), but for HepG2 cells. WT Ctrl n=1226 mitochondria from N=36 cells; WT ATP N=27, n=1301; *Efhd1*^-/-^ Ctrl N=25, n=972; *Efhd1*^-/-^ ATP N=27, n=1068. **I.** Mean mitochondrial size. WT Ctrl: n=1226 mitochondria from N=36 cells; *Efhd1*^-/-^ Ctrl: N=25, n=972; EFHD1-HA: N=41, n=530. **J, K.** Fission events counted after vehicle (PBS, ctrl) or 1mM ATP. Hepatocytes: MitoTracker label; WT Ctrl n=15; WT ATP n=9; *Efhd1*^-/-^ Ctrl n=19; *Efhd1*^-/-^ ATP n=12. HepG2: mito-mGold2s label; WT Ctrl n=22; WT ATP n=29; *EFHD1*^-/-^ Ctrl n=19; *EFHD1*^-/-^ ATP n=20. **L, M.** Representative images (L) and summary of mitochondrial length (M). Top, boxed insets in (L) at higher magnification. WT n=14; *Efhd1*^-/-^ n=9. Bars: mean ± SEM.

That mitochondrial remodeling was associated with ER Ca^2+^ release was established >20 years ago, but its significance, mechanism, and sensor remain elusive(32). Therefore, we tested whether EFHD1 mediates Ca^2+^-dependent mitochondrial remodeling, by inositol triphosphate receptor-mediated triggering ER Ca^2+^ release with 1 mM extracellular ATP (**Fig. 1E-F, S9D-E**) (33). This led to a marked reduction in size in WT but not *Efhd1^-/-^* mitochondria (**Fig. 2A-B, 2G-H**). Next, we focused on mitochondrial fission, a well-defined outcome of stable ERMCS formation(34). We found that deleting EFHD1 in both hepatocytes and HepG2 cells prevented the increase in fission events following ATP (**Fig. 2J-K**). Moreover, fission was also reduced at baseline in EFHD1-deficient cells. To confirm Ca^2+^ dependence, we permeabilized plasma membranes with a low concentration of digitonin, leaving mitochondria intact, and added a 100 µM Ca^2+^ bolus. In WT cells, mitochondria both rounded up and fissioned, an effect substantially blunted in *Efhd1^-/-^* cells (**Fig. 2L-M**). Finally, this effect was linked to nutritional state. Adding glucose and palmitate to starved hepatocytes remodeled WT but not *Efhd1^-/-^* mitochondria (**Fig. S3K**). Taken together, these results establish EFHD1 as a necessary transducer of ER Ca^2+^ release into a mitochondrial remodeling signal.

### EFHD1-dependent actin bundling integrates spatial proximity with a temporal Ca^2+^ signal to stabilize ERMCS

Next, we investigated how EFHD1 drives Ca^2+^-dependent remodeling. When ERMCS form, actin filaments are thought to assemble by an interaction between the ER-associated protein INF2 and mitochondria-associated Spire1C (**Fig. 3A**)(35, 36). For subsequent fission, the cytoplasmic GTPase DRP1 is recruited by adaptors, and constricts the mitochondrion until it divides. Prior studies have suggested Ca^2+^ can drive INF2-dependent actin polymerization, aid in the recruitment of DRP1 to fission sites during cell injury, and/or drive energetics within the mitochondrial matrix (37–40). Thus, how this Ca^2+^ signal drives remodeling remains undefined.

**Figure 3.**
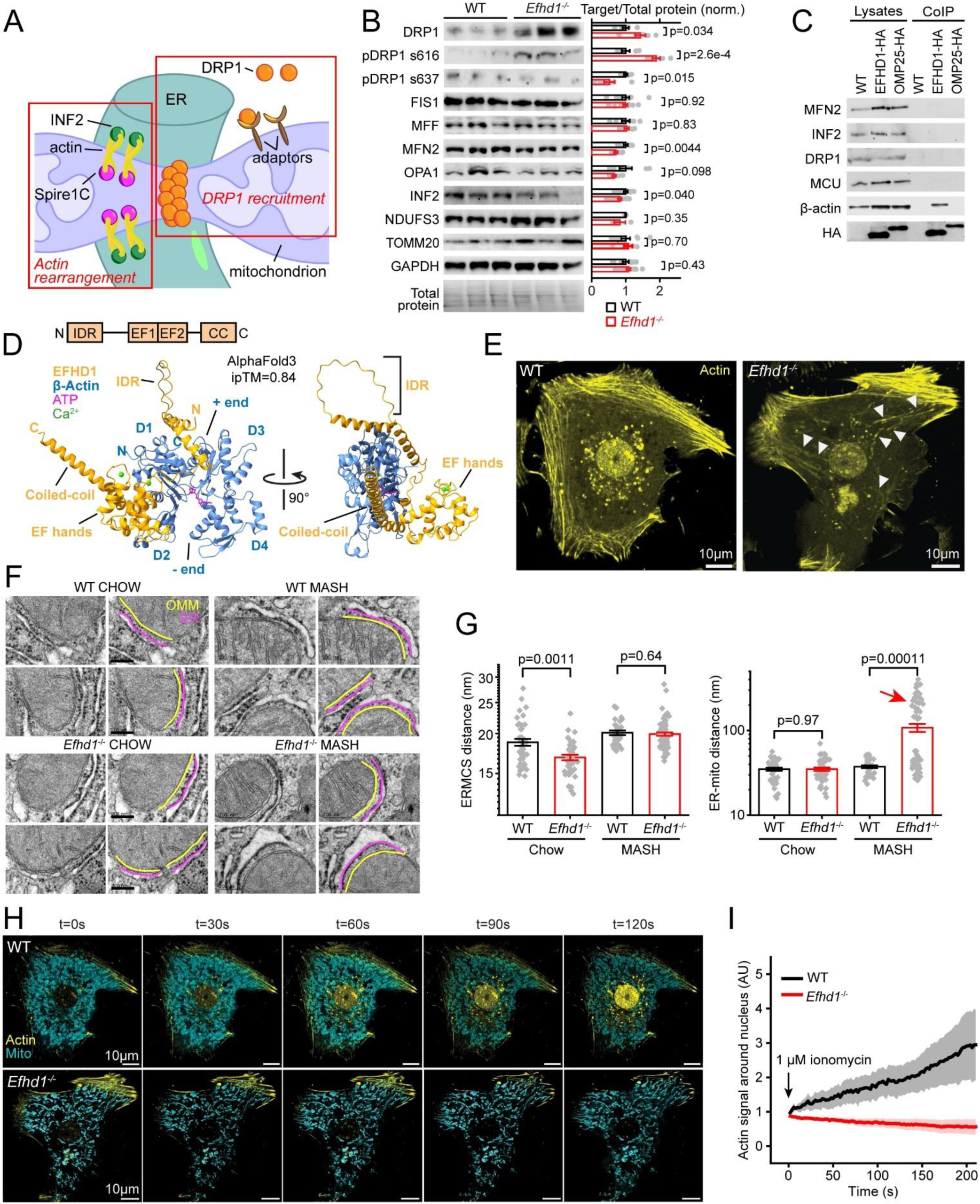
EFHD1 triggers Ca^2+^-dependent ERMCS stabilization. **A.** Important events to establish ERMCS (actin rearrangements) and initiate fission (DRP1 recruitment). **B.** Liver protein western blot and summary (n=6 per genotype, normal chow). **C.** Co-immunoprecipitation of HepG2 cells, untransfected (WT), transfected with EFHD1-HA or OMP25-GFP-HA (outer membrane protein). Representative of 4 experiments. **D.** High-confidence EFHD1:β-actin AlphaFold3 interaction. EFHD1 domains are shown above. ATP and Ca^2+^ are placed correctly within the EFHD1 EF hands and actin nucleotide binding pocket, respectively. **E.** Representative SiR-Actin-stained hepatocytes. Arrows, cytoplasmic stress fibers. **F.** Representative TEM for ERMCS. Left, plain images. Right, OMM (yellow) and ER membrane (magenta) at ERMCS highlighted. **G.** ER-mitochondrial distance at ERMCS (left) or any close association (<500 nm) on TEMs. Arrow points to a new population in MASH *Efhd1^-/-^* hepatocytes^..^ N=3 mice for each condition where n=135 micrographs (WT CHOW); n=150 (*Efhd1*^-/-^ CHOW); n=90 (WT GAN); n=92 (*Efhd1*^-/-^ GAN). **H, I.** Representative images (H) and summary (I), peri-nuclear SiR-actin signal in chow-fed hepatocytes. WT: N=3 mice and n=11 cells. *Efhd1^-/-^* : N=3, n=8. Bars: mean ± SEM.

We began by testing whether the Ca^2+^ signal for remodeling acts at the OMM or within mitochondria(37). In all the experiments described so far, we had incubated cells with 1 µM Ru360, a blocker of the mitochondrial Ca^2+^ uniporter, the channel conducting Ca^2+^ into the matrix, implying that remodeling did not require Ca^2+^ uptake. To confirm this further, we silenced the main uniporter subunit MCU with short-hairpin RNA (**Fig. S4A-B**)(41), finding again that depleting MCU had no effect on ATP-induced fission events. Because prior work suggested EFHD1 can bind the mitochondrial Ca^2+^ uniporter, we also examined whether EFHD1 modulates hepatocyte mitochondrial Ca²⁺ entry(30). Across assays of uniporter protein levels, mitochondrial Ca^2+^ uptake, and mitoplast patch-clamp measurements, we detected no differences between wild-type and *Efhd1^-/-^* hepatocytes (**Fig. S4C-K**). The imaging assays with ATP (**Fig. S4E-G**) also confirmed that differences in remodeling seen in **Fig. 2** were not due to aberrant ER Ca^2+^ release. Thus, the Ca^2+^ signal driving EFHD1-dependent mitochondrial remodeling localizes to the OMM.

Next, we tested the hepatic expression levels of proteins involved in mitochondrial dynamics (**Fig. 3B**). The most prominent change was a marked increase in DRP1 in *Efhd1*^-/-^. This was not due to changes in mitochondrial content, as mitochondrial markers NDUFS3 and TOMM20 were unchanged. This result is counterintuitive, since increased DRP1 would be expected to produce excessive fission in *Efhd1^-/-^*, rather than the elongated mitochondria we observe. An explanation might be that DRP1 is not migrating to the OMM, since prior studies established that, during cell damage, Ca^2+^ promotes DRP1 localization to the OMM by enhancing Ser-616 phosphorylation and Ser-627 dephosphorylation(39, 40). Consequently, we examined DRP1 OMM recruitment. Unexpectedly, DRP1 Ser-616 phosphorylation was increased, while Ser-637 phosphorylation was reduced in *Efhd1^-/-^* livers, suggesting DRP1 was hyperactive (**Fig. 3B**). Moreover, DRP1 puncta localized to mitochondria in both WT and *Efhd1^-/-^* cells, further confirming no defect in its activity (**Fig. S5A**). Therefore, EFHD1 is not directly affecting mitochondrial fission. Rather, the increased expression of hyperactive DRP1 likely contributes to the low residual fission still present in *Efhd1^-/-^* cells, and along with the decreased expression of fusion proteins MFN2 and OPA1 (**Fig. 3B**), forms a compensatory response for an impairment at an earlier step in ERMCS formation.

Since EFHD1 is not directly involved in DRP1 activation or downstream fission, we examined Ca^2+^-dependent actin rearrangements at ERMCS(35). In fact, EFHD1 and its homolog EFHD2 are known to bundle actin filaments in the presence of Ca^2+^(42, 43). Moreover, in recent protein-protein interaction compendia, EFHD1 is annotated as an actin-binding protein, which we confirmed via coimmunoprecipitation (**Fig. 3C, S5B**)(29). Expression of INF2, which polymerizes actin at ERMCS but does not itself bind Ca^2+^, was also reduced in *Efhd1^-/-^*, though it did not co-immunoprecipitate with EFHD1 (**Fig. 3B-C**). Next, we modeled binding via AlphaFold3, which predicted a high-confidence interaction between EFHD1 and β-actin (**Fig. 3D, S5C**)(44). In the predicted structure, EFHD1 was primarily bound between domains 1 and 2 of actin, leaving free the plus- and minus-ends, similar to other actin-bundling proteins (**Fig. S5D**)(45, 46). The greatest uncertainty in the prediction was the localization of the EFHD1 initial helical domain, with several models suggesting it could interact with the actin plus-end, potentially regulating actin polymerization (**Fig. S5C**).

We then investigated whether loss of EFHD1 would alter actin networks, ERMCS, and Ca^2+^-dependent actin remodeling. First, whereas control cells had diffuse cytoplasmic actin staining, cells lacking EFHD1 displayed prominent cytoplasmic actin stress fibers, away from peripheral cortical networks (**Fig. 3E, S5E**). Second, we examined ERMCS, where mitochondria and ER are <30 nm apart (**Fig. 3F-G**)(47). We performed two analyses, measuring either the distance at ERMCS below the 30 nm threshold, or distances whenever there were stretches of ER and OMM <500 nm apart. Notably, in animals fed a chow diet, ERMCS were narrower for *Efhd1*^-/-^hepatocytes compared to WT, suggesting ER and mitochondria needed to be closer to establish robust contacts. In addition, while ERMCS distances were stable between WT animals fed either a normal or MASH diet, ER-mitochondrial distances were widened and irregular in *Efhd1^-/-^*hepatocytes in MASH, including distinct populations with large separations not present at baseline (**Fig. 3G**). The failure to maintain ERMCS integrity during MASH indicated that EFHD1 is required for stabilizing these contacts(48). Finally, we directly measured the Ca^2+^- and INF2-dependent cellular actin response using a well-established protocol(37, 38). Here, ionomycin increases cytoplasmic Ca^2+^ to drive actin polymerization, visualized as a peri-nuclear actin ring in WT hepatocytes but entirely absent in *Efhd1*^-/-^ hepatocytes (**Fig. 3H, I**). We conclude from these experiments that EFHD1 is a spatiotemporal coincidence detector that helps create stable ERMCS. ER Ca^2+^ release is the trigger for EFHD1 to bundle actin from opposite membranes, efficiently tethering ER to mitochondria (**Fig. S6**). In the absence of EFHD1, ERMCS tethering requires a direct but unstable INF2-Spire1C interaction, which produces narrower ERMCS that are more prone to disruption.

### EFHD1 ablation reduces hepatocyte inflammation and the integrated stress response

Next, we used a multi-omic approach to define the link connecting remodeling to MASH. Here, we primarily used male mice, since differences in their liver phenotypes were strongest. We performed RNA-seq of total RNA and label-free proteomic analysis of entire livers of WT and *Efhd1^-/-^* mice fed either normal or MASH diets (**Fig. S7A-D, Table S1-2**). MASH diet fed animals had more pronounced differences between WT and *Efhd1^-/-^* compared to normal chow, but in both diets the most notable effect of EFHD1 deletion was a decrease in interferon inflammatory pathways (**Fig. 4A-B**). We also performed label-free proteomic analysis of entire livers (**Fig. S7E-G, Table S3-5**). We analyzed this data in three ways: looking at changes between WT and *Efhd1^-/-^* for both diet conditions (**Fig. 4C, E, F, H**), but also examining changes in WT between normal and MASH diets (**Fig. 4D, G**). This additional analysis allowed us to examine which specific pathways dysregulated by MASH are rescued by EFHD1 ablation (**Fig. 4I-L**). Using REACTOME pathway analysis, the most prominent changes caused by MASH versus normal diets were increased fibrosis, decreased mitochondrial metabolism, and complement pathway activation (**Fig. 4D, G**). For *Efhd1^-/-^* livers, we again found decreases in inflammatory and fibrosis pathways and improved mitochondrial metabolism.

**Figure 4.**
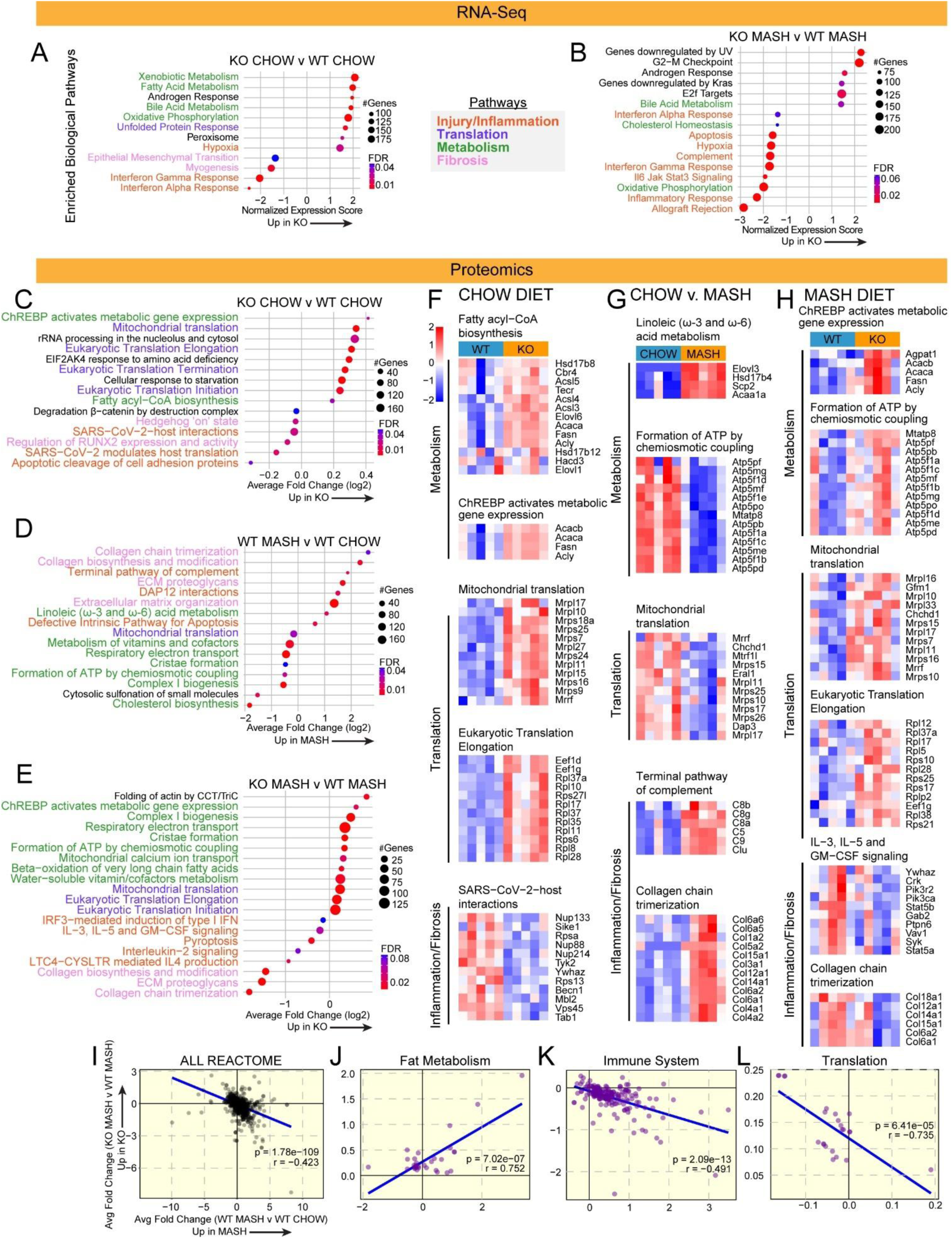
Transcriptomic and proteomic analysis of EFHD1 ablation. A, **B.** Selected Hallmark pathways differentially regulated in whole liver RNA-seq of *Efhd1^-/-^* versus WT mice. FDR, false discovery rate. **C-E.** Selected REACTOME pathways differentially regulated in whole liver proteomics. **F-H.** Heat maps of selected proteins from each indicated REACTOME pathway. **I-L.** Fold change for WT MASH versus WT normal chow (x axis) graphed against the fold change for *Efhd1^-/-^* MASH versus WT MASH (y axis) for parental REACTOME pathways (I), fat metabolism pathways (J), immune system pathways (K), and translation pathways (L). Changes in MASH (relative to normal chow) reversed in *Efhd1^-/-^* is visible as a negative slope. For (K), increases in immune pathways during WT MASH are decreased in *Efhd1^-/-^* MASH, whereas for (L), decreases in translation during WT MASH are increased in *Efhd1^-/-^* MASH. RNA-seq, chow: WT n=5, *Efhd1^-/-^* n=5. MASH: WT n=4, *Efhd1^-/-^* n=5. Proteomics, chow: WT n=5, *Efhd1^-/-^* n=5. MASH: WT n=5, *Efhd1^-/-^* n=5.

To examine such changes further, we plotted the fold change for each pathway in the “WT MASH versus WT normal” comparison against the fold change in that pathway in the “*Efhd1^-/-^* MASH versus WT MASH” comparison (**Fig. 4I-L**). A negative slope shows that pathways upregulated by MASH in WT animals are downregulated in *Efhd1^-/-^* livers, and vice versa, confirming that pathways dysregulated in MASH tend to recover in the *Efhd1^-/-^* animals globally (**Fig. 4I**). Focusing on specific parental REACTOME groups of interest, we found that Fat Metabolism was one cluster that did not show a significant relationship (**Fig. 4J**). In striking contrast, broad increases in Immune System pathways during MASH were downregulated in *Efhd1^-/-^* animals (**Fig. 4K**), consistent with the histopathological phenotypes.

An unexpected and intriguing set of MASH-dysregulated pathways rescued by EFHD1 ablation were those related to translation (**Fig. 4E, H, L**). Global decreases in mRNA translation are a hallmark of the integrated stress response (ISR). In the ISR, phosphorylation of the elongation initiation factor 2A (EIF2A) globally downregulates translation, with preferential expression of ATF4-dependent genes that promote cellular recovery(49). Whereas temporary ISR activation is protective, prolonged activation promotes cell death. The ISR and a closely-related mechanism, the unfolded protein response (UPR), are activated in the liver by ER lipotoxicity(50). The ISR can also be activated by severe mitochondrial dysfunction, double-stranded RNA associated with viral infection, or starvation(49). In our proteomic dataset, translation was decreased in MASH, consistent with ISR activation, and rescued by ablating EFHD1, suggesting reductions in ISR in *Efhd1^-/-^* mice. Taken together, these data are consistent with histopathological and GWAS data suggesting that EFHD1 inhibition primarily prevents the hepatocyte injury leading to inflammation and fibrosis, with much less pronounced effects on lipid metabolism. Furthermore, these data point towards the ISR as a possible mechanism linking mitochondrial function to hepatocyte injury.

### PKR activation is via pathological release of mitochondrial double-stranded RNA

To investigate the ISR further, we assessed the expression and phosphorylation state of EIF2α, upstream kinases, and ATF4 target genes(51). As expected, in MASH we found substantial upregulation of EIF2α phosphorylation, PERK expression and phosphorylation, and ATF4 targets (**Fig. 5A-B**, **Fig. S8A**). In contrast, compared to WT, *Efhd1^-/-^* livers on a MASH diet showed reduced ISR (**Fig. 5C-D, S8B**).

**Figure 5.**
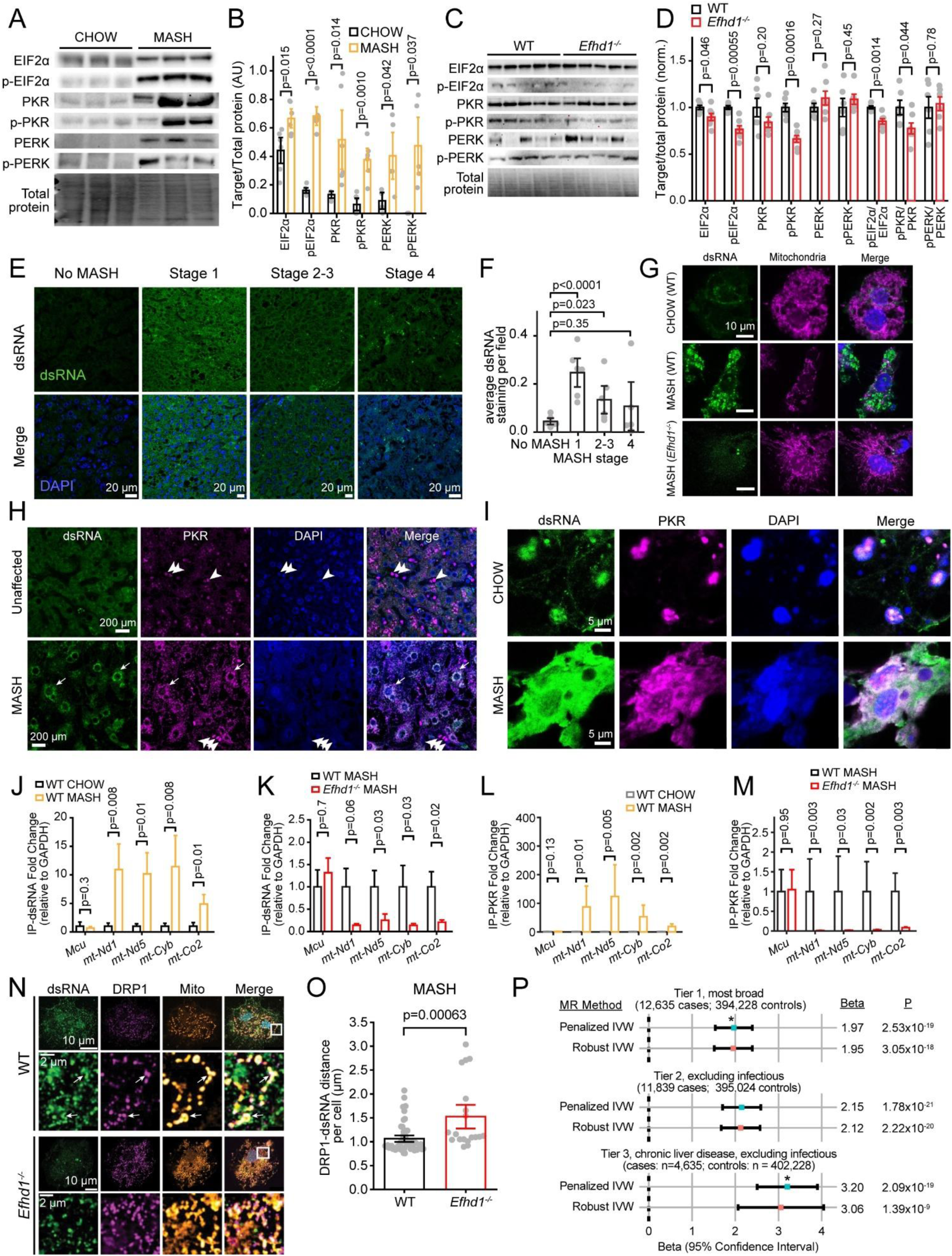
Excessive ERMCS stabilization triggers pathological mt-dsRNA release and a PKR-dependent integrated stress response in MASH. A, **B.** Western blot (A) and summary (B) EIF2a: Eukaryotic translation initiation factor 2A; p-EIF2a: Phosphorylated EIF2a; PKR: Protein kinase R; p-PKR: Phosphorylated PKR; PERK: protein kinase R (PKR)-like endoplasmic reticulum kinase/Eukaryotic translation initiation factor 2-alpha kinase 3 (EIF2AK3); p-PERK: Phosphorylated PERK. Chow n=6, MASH n=5. **C, D.** MASH diet western blot (C) and band quantification (D). WT n=7, *Efhd1^-/-^* n=7. **E, F.** Representative images (E) and quantification summary (F) of dsRNA immunohistochemistry in human liver (Stage 0, 1, 2-3 n=6, Stage 4 n=5). **G.** Isolated mouse hepatocyte immunohistochemistry. **H.** Representative high-contrast immunohistochemistry of human livers. Arrowheads, PKR in dense non-hepatocyte nuclei; arrows, dsRNA- and PKR-stained cytoplasmic signal. **I.** Representative images, isolated mouse hepatocytes. **J-M.** Quantitative RT-PCR following immunoprecipitation of dsRNA (J, K) or PKR (L, M) from isolated mouse hepatocytes. dsRNA: WT chow n=8 ; WT MASH n=6 ; *Efhd1^-/-^* MASH n=8. PKR WT chow n=5; WT MASH n=5; *Efhd1^-/-^* MASH n=7. **N.** Representative immunocytochemistry of isolated hepatocytes from MASH diet mice. Because less dsRNA was evident in *Efhd1^-/-^* hepatocytes, image acquisition settings for these were adjusted to maximize dsRNA signal. Bottom, inset at higher magnification. **O.** Average dsRNA-DRP1 distance per hepatocyte (WT n=49, *Efhd1^-/-^* n=19, N=3 mice per condition). **P.** Penalized and robust inverse variance-weighted MR analyses support a causal relationship between human *PKR* (*EIF2AK2*) and liver disease, using AST as the exposure. *, two distributions with non-overlapping 95% confidence intervals.

Our next goal was to define the damage mechanism activating the ISR. Notably, a well-established inflammation trigger in other cell types, release of mitochondrial DNA (mtDNA), fails to activate an inflammatory response in hepatocytes because they lack STING expression(52–54). Another mitochondrial damage-associated signal may arise from improper processing of mitochondrial RNA (mt-RNA). Normally, this RNA is transcribed as two separate but complementary strands, each cleaved to release specific transcripts. During mitochondrial dysfunction, mt-RNA is inadequately processed, and can hybridize and leak into the cytoplasm as a long dsRNA(55, 56). Within the cytoplasm, mt-dsRNA may be recognized by the dsRNA-sensor PKR (Protein kinase RNA-activated, *EIF2AK2* gene), which can trigger the ISR independently of PERK. Notably, hepatic PKR typically senses dsRNA from viruses such as hepatitis C, activating ISR to prevent viral replication(57). Thus, mt-dsRNA-dependent PKR activation may represent a maladaptive response.

The possibility that PKR might link mt-dsRNA to the ISR during MASH was interesting for additional reasons. Notably, we saw marked increases in PKR and phospho-PKR in MASH diet-fed mice, comparable to the increases in PERK (**Fig. 5A-B, S8A**). Moreover, whereas PERK signaling was not affected by EFHD1 deletion, we saw decreases in phospho-PKR in MASH-fed *Efhd1^-/-^* mice relative to WT (**Fig. 5C-D**). In contrast, two other dsRNA sensors involved in innate immunity, RIGI and MDA5, were not substantially altered (**Fig. S8C-D**). Finally, prior studies investigating overnutrition found that PKR deletion reduced organismal inflammation, but disagreed on whether lipid metabolism was affected, and failed to identify a trigger for PKR activation(58, 59). This pattern of protection from inflammation with variable effects on metabolism mimics the phenotype of EFHD1 ablation.

Therefore, to determine whether mt-dsRNA triggered the ISR in MASH, we stained human liver tissue sections for dsRNA(55). This revealed a clear increase in dsRNA signal during early MASH (**Fig. 5E-F**). In later stages, the dsRNA signal became more heterogenous but remained elevated. Widespread cytoplasmic dsRNA staining was also evident in isolated hepatocytes from WT mice fed a MASH diet (**Fig. 5G, S8E-F**)(60). In contrast, there was minimal dsRNA staining in both chow-fed WT mice and in *Efhd1^-/-^* mice fed a MASH diet. We then co-stained for both dsRNA and PKR in human and mouse livers (**Fig. 5H-I, S8G-H**). In healthy tissue, PKR localized to condensed nuclei in non-hepatocyte cells, likely leukocytes(61). Conversely, in MASH, PKR staining was now also present in hepatocytes and overlapped the cytoplasmic dsRNA stain. Therefore, in both humans and mice, overnutrition triggers the cytoplasmic release of endogenous dsRNAs, where they are sensed by PKR.

Deletion of EFHD1 prevented the pathologic increase in cytoplasmic dsRNA, suggesting mitochondrial origin. To confirm this, we immunoprecipitated either dsRNA or PKR from isolated hepatocytes and quantified bound mitochondrial RNA via quantitative reverse transcription polymerase chain reaction(56). Compared to a nucleus-encoded mitochondrial gene (*Mcu*), mt-RNAs were profoundly enriched in the dsRNA- or PKR-immunoprecipitated fraction in MASH-fed WT mice (**Fig. 5J-M**). However, in *Efhd1^-/-^* mice fed a MASH diet, the degree of mt-dsRNA enrichment was much reduced (**Fig. 5K, M**). Finally, we found that dsRNA localized near DRP1 and mitochondria in MASH livers, linking mt-dsRNA release to excessive constriction (**Fig. 5N**). In *Efhd1^-/-^* MASH hepatocytes, though dsRNA could be visualized when increasing image acquisition settings, the distance to DRP1 was increased (**Fig. 5O**). Taken together, our data suggest that EFHD1 upregulation during MASH drives ERMCS hyperstability and mitochondrial fragmentation, causing the release of mt-dsRNA and the subsequent activation of a maladaptive PKR-dependent ISR.

### Mendelian randomization studies support a causal relationship between human *PKR (EIF2AK2)* and liver disease

To investigate whether PKR produces a similar phenotype to EFHD1 in humans, we first queried the Common Metabolic Diseases Knowledge Portal(62). As with *EFHD1*, variation in the *PKR (EIF2AK2* gene) locus was associated with serum liver enzymes but not triglycerides, liver fat, or other lipid metabolism parameters (**Fig. S8I**). Furthermore, analyzing the UK Biobank, we identified statistically significant associations (p-value < 8.87×10^-6^) between 35 genetic variants spanning a 105.8 kilobase region at the *EIF2AK2* locus and serum AST levels (**Fig. S8J, Table S6**). Significant eQTLs (p-value < 8.23×10^-4^) for *EIF2AK2* expression in liver were observed at 34 of these AST-associated variants (**Fig. S8K**)(63). 21 of these variants were also predicted by RegulomeDB rank and model prediction scores to have a high likelihood of being functional(64).

To further support a causal association, Mendelian randomization (MR) was performed on these 35 variants at the *EIF2AK2* locus, using AST as the exposure and liver disease diagnosis as the outcome. MR is a causal inference method that uses genetic variants as instrumental variables to test the causal effect of that trait on a disease outcome. By leveraging the random assortment of alleles at conception, MR minimizes confounding and reverse causation, providing unbiased evidence for a causal link in humans. We ran three MR analysis that varied by liver diagnosis inclusion (**Fig. 5P, Table S7-11**). Tier 1 was the broadest, including diagnoses associated with acute or chronic liver disease, malignancies, and infectious causes. In tier 2, we excluded infectious causes, including viral hepatitides. Tier 3 was the most restrictive, including only codes associated with chronic liver disease and excluding infectious causes. MR analysis for all three outcomes suggested a causal association between AST-associated variants and liver disease. Strikingly, the magnitude of the penalized IVW estimate for Tier 3 was significantly larger than for Tier 1, suggesting a stronger effect of PKR on chronic liver disease not due to viral infection. Overall, this putative causal relationship reinforces a pathway linking aberrant mt-dsRNA release to hepatocyte injury.

### Liver-specific EFHD1 inhibition in humans and mice is hepatoprotective

Our final goal was to establish whether liver-specific EFHD1 inhibition could be hepatoprotective. As expected, in mice with liver-specific EFHD1 ablation (*Efhd1^hKO^*), hepatocytes had elongated mitochondria resistant to Ca^2+^-induced fission (**Fig. 6A-D, S9A-B**). Then, we injected 8–10-week-old male *Efhd1^hKO^*or *Alb*-Cre control mice with carbon tetrachloride (2 µL/g CCl_4_, 3x week) intraperitoneally for 6 weeks, a chemical liver injury model that produces marked fibrosis rather than steatosis(65). Again, *Efhd1^hKO^* mice had reduced serum liver enzymes and fibrosis (**Fig. 6E-I**). This result revealed that EFHD1 mediates a core injury response independent from the initial insult.

**Figure 6.**
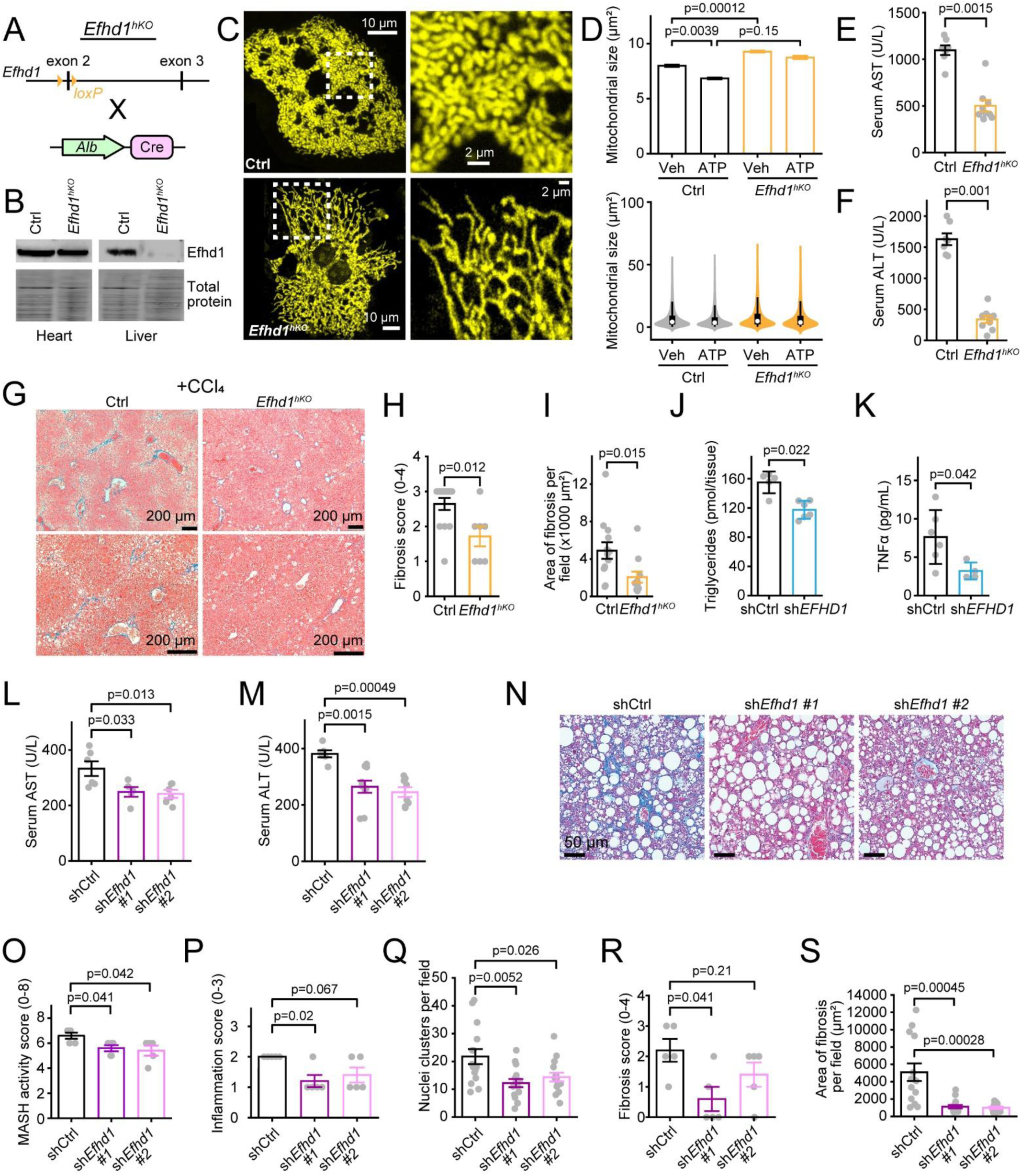
Liver-specific EFHD1 inhibition reduces hepatocyte injury. **A.** Hepatocyte-specific *Efhd1* deletion via frameshift and early termination. **B.** Western blot. **C.** MitoTracker-stained isolated mouse hepatocytes. Right, boxed insets at higher magnification. **D.** Automated iLastik analysis. Bar chart (top) shows mean effects; violin plot (bottom) shows full distribution. WT Ctrl n=12055 mitochondria from N=6 mice; WT ATP N=4, n=19498; *Efhd1^hKO^* Ctrl N=3, n=24890; *Efhd1^hKO^* ATP N=3, n=21088. **E-I.** Assays following 4 weeks CCl_4_ injection. **E, F.** Serum liver enzymes. **G.** Liver Masson’s trichrome staining. Fibrosis, blue. **H.** Histopathological fibrosis score. **I.** Fibrotic area from liver micrographs. **J-K,** Human organoid (J) triglyceride (shCtrl n=5, sh*EFHD1* n=6), and (K) TNFα production (shCtrl n=6, sh*EFHD1* n=4). **L-S.** Assays on MASH diet and AAV8 treatment. **L, M.** Serum liver enzymes. **N.** Liver micrographs of Masson’s trichrome staining. **O.** MASH activity score. **P.** Histopathological inflammation score. **Q.** Liver leukocyte cluster counts. **R.** Histopathological fibrosis score. **S.** Fibrotic area measured from liver micrographs. B-I, L-S used male mice. Clinical histopathological scoring in H, O, P, R was assessed in blinded fashion by liver pathologist (K.J.E.).For E-F, WT n=7; *Efhd1*^hKO^ n=9. For H, WT n=14; *Efhd1*^hKO^ n=7. For I, WT n=13; *Efhd1*^hKO^ n=11. For N, shCtrl n=6; sh*Efhd1#1* n=5; sh*Efhd1#2* n=6. For O, shCtrl n=6; sh*Efhd1#1* n=10; sh*Efhd1#2* n=8. For O, P, R, shCtrl n=5; sh*Efhd1#1* n=5; sh*Efhd1#2* n=5. For Q, S, shCtrl n=15; sh*Efhd1#1* n=15; sh*Efhd1#2* n=15. Bar: mean ± SEM.

Next, we investigated whether acute inhibition of EFHD1 could confer similar protection. First, we examined human 3D liver microtissues that were transduced before aggregation with either a non-targeting control or *EFHD1-*targeting short hairpin RNA (∼85% knockdown of transcripts), packaged in adeno-associated virus (AAV-DJ, **Fig. S9C-D**)(66). All viruses also encoded GFP, allowing us to confirm robust liver transduction. Both TNFα secretion and microtissue triglyceride content were reduced, suggesting that acutely targeting EFHD1 could benefit inflammation and injury in these human models (**Fig. 6J-K**).

Second, 6-week-old male mice were fed a MASH diet for 18 weeks before a single injection of *Efhd1-*targeting short hairpin RNA, or a non-targeting control, packaged in a hepatotrophic adeno-associated virus serotype (AAV8) (**Fig. S9E-F**). Livers were assayed 12 weeks later. This protocol allowed us to observe the effects of *Efhd1* inhibition after MASH diet-induced injury(67). We used two independent shRNAs targeting *Efhd1,* with sh*Efhd1*#1 producing greater inhibition than sh*Efhd1*#2 (**Fig. S9G-J**).

After *Efhd1* inhibition we again observed substantial decreases in liver enzymes, indicative of reduced liver injury (**Fig. 6L-M**). Unexpectedly, serum triglycerides as well as direct measures of lipid content, though not histopathological scores, also showed decreases, suggesting that acute EFHD1 inhibition led to downstream improvements in metabolism by preventing hepatocyte injury (**Fig. S9K-N**). Finally, both histological and direct scoring showed improvements in inflammation and fibrosis (**Fig. 5N-S, S9O**). In all these analyses, sh*Efhd1*#1 tended to produce stronger effects than sh*Efhd1*#2, consistent with a dose-response relationship between the degree of *Efhd1* inhibition and hepatic protection. Taken together, inhibiting EFHD1 is hepatoprotective across multiple human and mouse models of liver disease.

## DISCUSSION

Inter-organellar communication is essential for cellular homeostasis, but the fundamental principles governing the establishment of these contact sites remain poorly understood. Here, we identify EFHD1 as a factor that creates stable ERMCS by transducing a timing signal, ER Ca^2+^ release, into mechanical stability via actin bundling. A coincidence detection system integrating spatial proximity with a temporal cue provides several advantages over the current model of tethering driven by constitutive membrane–membrane interactions (**Fig. S6**). First, such interactions may be relatively inefficient, since these two proteins reside on different membranes. In support of this, ER and mitochondria in *Efhd1^-/-^* hepatocytes needed closer contact to establish a connection, evident in narrower ERMCS distances at baseline (**Fig. 3G**). Second, by requiring Ca²⁺ binding, EFHD1 ensures that contact formation is event-driven, occurring only at locations where the ER signals to mitochondria, and thereby preventing inappropriate or excessive contact during organelle motility in a crowded cytoplasm. Third, because EFHD1 transduces this signal specifically by actin bundling, it provides a way to generate mechanical stability at contact sites, evident in the disruption of ERMCS during metabolic stress in *Efhd1^-/-^* hepatocytes. Taken together, our findings reveal the molecular logic for robust ERMCS stabilization.

EFHD1 inhibition did not substantially affect organismal lipid metabolism or energy balance. Though we found that acute EFHD1 inhibition improved serum triglycerides, and there was less steatosis in *Efhd1^-/-^* mice compared to controls on a normal diet, this difference disappeared in the hepatotoxic diets. Moreover, across different cell types, there is no consistent effect on metabolism after EFHD1 inhibition(12–15). Fatty acid synthesis genes were upregulated in liver organoids, but lipogenesis was unaffected in HepG2 cells(12, 13). Thus, the variability in these results suggest the beneficial effects from EFHD1 inhibition on metabolism may be indirect, because hepatocytes are less injured or have less suppressed translation.

In contrast, EFHD1-dependent ERMCS stabilization creates an intrinsic vulnerability that directly leads to hepatocyte injury across all the models and conditions. EFHD1 upregulation during metabolic stress leads to pathological contact hyperstability, excessive mitochondrial fragmentation, and mt-dsRNA leakage. In the liver, cytoplasmic dsRNA is typically encountered during viral infection, where PKR drives the ISR to prevent translation of viral RNA. Here, we establish that during MASH, pathological mt-dsRNA release causes maladaptive activation of this pathway (**Fig. S16P**). We establish a candidate causal relationship between PKR and liver injury in humans, a relationship that is surprisingly strongest for non-infectious chronic liver disease. Inhibiting EFHD1 protects the liver by selectively dampening this PKR-dependent ISR without disrupting PERK-driven responses. Because EFHD1 is a central node integrating Ca^2+^ signaling, organelle dynamics, and immune activation, targeting its activity provides a therapeutic strategy fundamentally different from metabolic modulation for the treatment of liver injury.

## METHODS

Detailed methods are available in the Supplemental Material.

## Supporting information

Combined supplementary file

## ACKNOWLEDGEMENTS

We thank Dr. Rajarshi Chakrabarti (Thomas Jefferson University) for guidance on actin assays. We thank Dr. David Clapham (Howard Hughes Medical Institute) for the kind gift of HepG2 cells, and Dr. Françoise St-Pierre (Baylor College of Medicine) for the kind gift of the mito-mGold2s construct. We thank Dr. Brian Dalley and staff at the Huntsman Cancer Institute High Throughput Genomics Core, Dr. Ying Li and staff at the University of Utah Metabolic Phenotyping Core, Dr. James Marvin and staff at the University of Utah Flow Cytometry Core, and staff at the University of Utah Electron Microscopy Core, and the University of Utah Research Histology Core. We thank the University of Utah Center for Metabolic Health for supporting access to UK Biobank.

