## Supplementary material for "Excessive Ca^2+^-dependent ER-mitochondrial contact stabilization by EFHD1 drives liver injury": Combined supplementary file

### SUPPLEMENTARY FIGURES

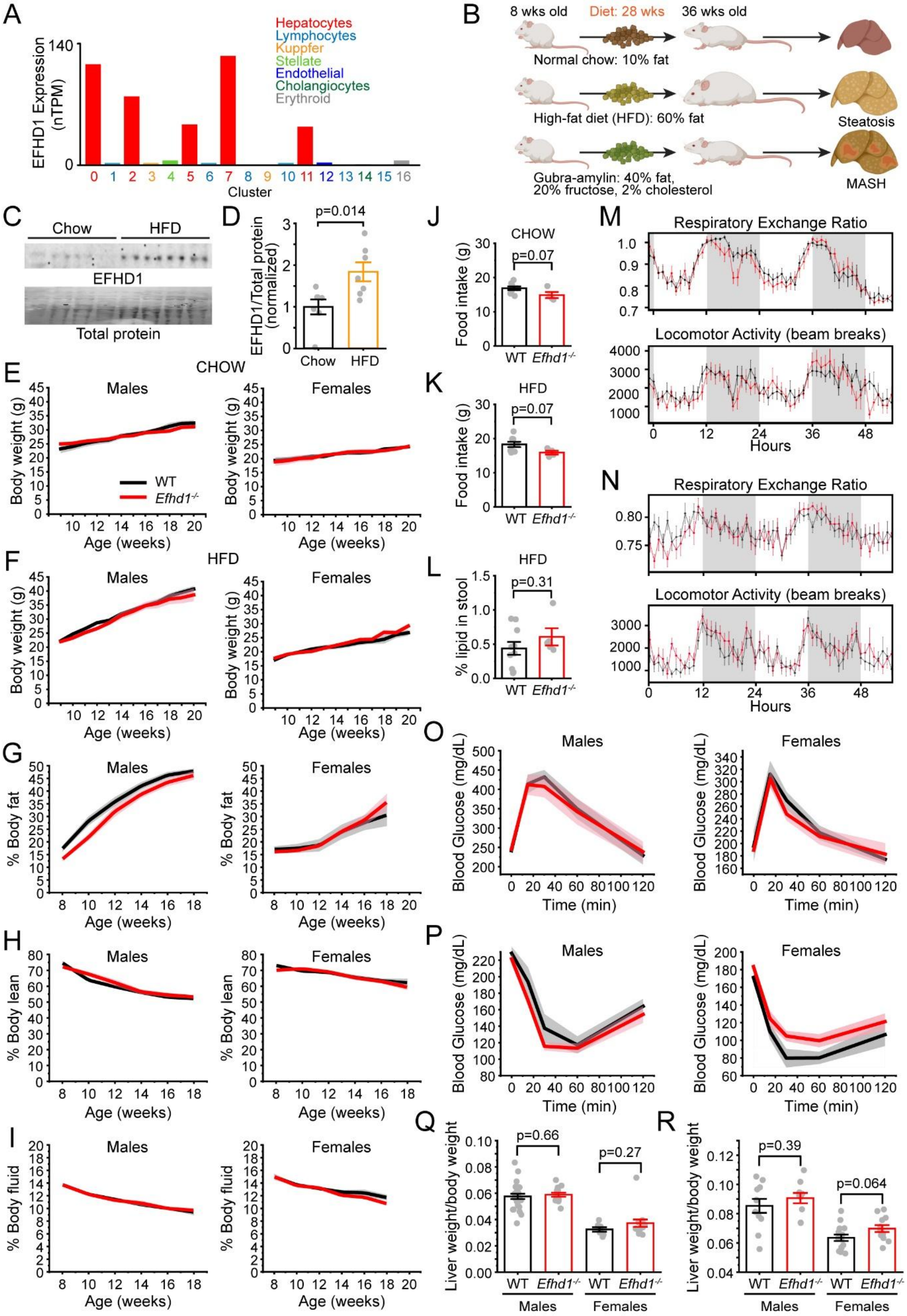

**Figure S1. Organismal energy balance is preserved in *Efhd1*$^{-/-}$ mice. A.** Human single-cell RNA-seq data reveals EFHD1 is preferentially expressed in hepatocytes (GSE115469). **B.** Timeline of dietary interventions. Created with BioRender. **C, D.** Western blot and band analysis for HFD-fed mice (D, n=7). **E, F.** Growth curves for normal chow and HFD over 20 weeks, showing no difference between WT and *Efhd1*$^{-/-}$ mice. Shaded grey and pink areas here and throughout represent SEM. (E) Normal Chow. WT: male n=7, female n=5; *Efhd1*$^{-/-}$: male n=5, female n=7. (F) MASH. WT: male n=12, female n=7; *Efhd1*$^{-/-}$: male n=5, female n=11. **G, H, I.** Body composition obtained for mice on 12 weeks HFD. (G) % body fat, (H) % body lean, and (I) and % body fluid. WT: male n=20, female n=11; *Efhd1*$^{-/-}$: male n=7, female n=15. **J, K.** Food intake over 5 days for normal chow and 20-week HFD-fed mice. WT, n=5; *Efhd1*$^{-/-}$, n=5. **L.** Stool lipid content in 20-week HFD-fed mice. WT, n=10; *Efhd1*$^{-/-}$, n=5. **M, N.** Metabolic cage assessment of respiratory exchange ratio (top) and locomotor activity (bottom) over 52 hours for mice fed normal chow (M; WT, n=5; *Efhd1*$^{-/-}$, n=5.) or 5 weeks HFD (N; WT, n=5; *Efhd1*$^{-/-}$, n=5.). **O, P.** Glucose (O) and insulin (P) tolerance tests for 12 weeks HFD-fed mice. WT: males n=8, females n=10; *Efhd1*$^{-/-}$: males n=8, females n=5. **Q, R.** Liver to body weight ratios for mice fed normal chow (Q, WT: males n=25, females n=7; *Efhd1*$^{-/-}$: males n=14, females n=14.) or HFD (R, WT: males n=10, females n=8; *Efhd1*$^{-/-}$: males n=14, females n=11). Bar: mean ± SEM.

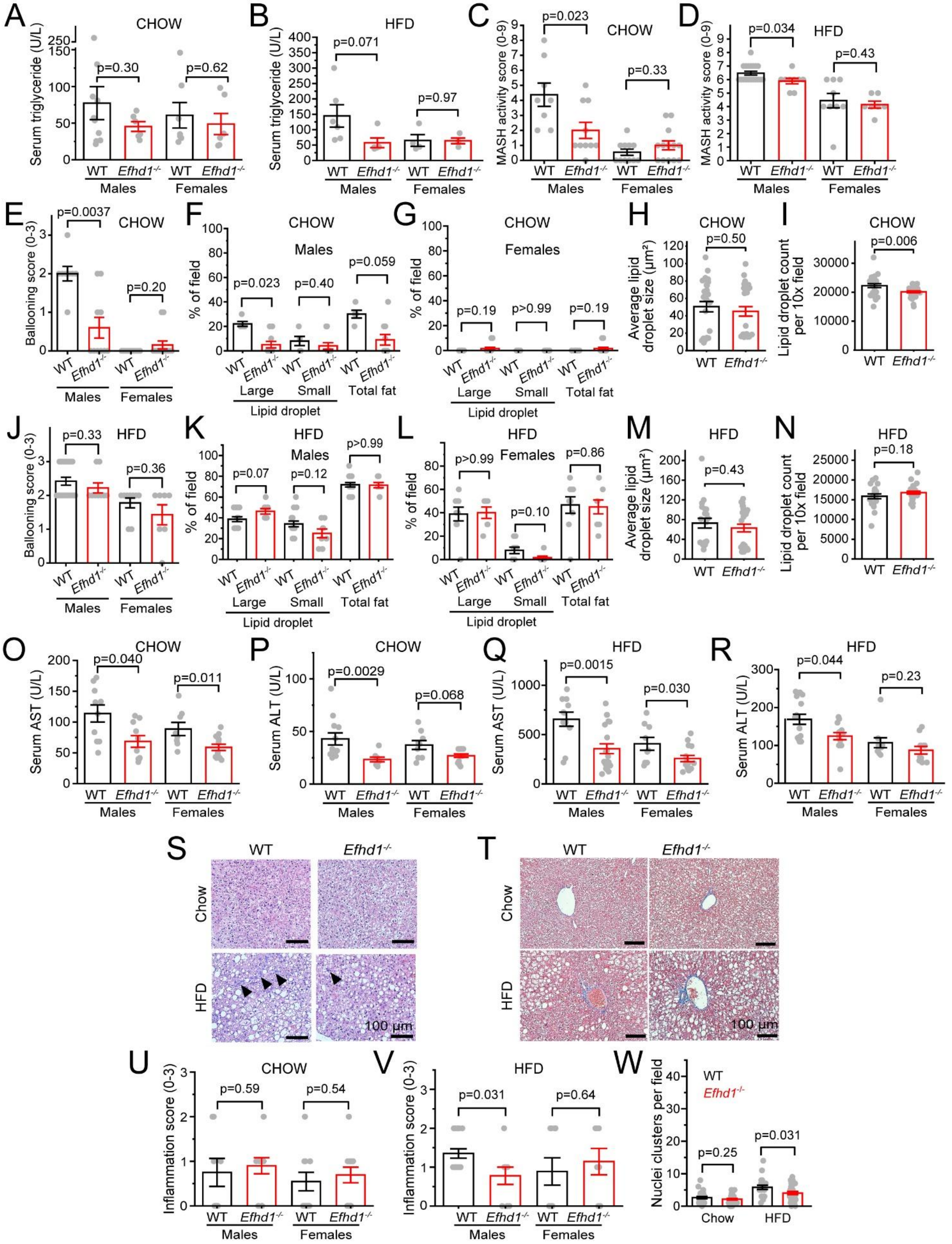
A
CHOW
Serum triglyceride (U/L)
p=0.30
p=0.62
WT
Efhd1-/-
Males
Females
B
HFD
Serum triglyceride (U/L)
p=0.071
p=0.97
C
CHOW
MASH activity score (0-9)
p=0.023
p=0.33
D
HFD
MASH activity score (0-9)
p=0.034
p=0.43
E
CHOW
Ballooning score (0-3)
p=0.0037
p=0.20
F
CHOW
Males
% of field
p=0.023
p=0.40
p=0.059
Large
Small
Total fat
Lipid droplet
G
CHOW
Females
% of field
p=0.19
p>0.99
p=0.19
H
CHOW
Average lipid droplet size (µm²)
p=0.50
I
CHOW
Lipid droplet count per 10x field
p=0.006
J
HFD
Ballooning score (0-3)
p=0.33
p=0.36
K
HFD
Males
% of field
p=0.07
p=0.12
p>0.99
L
HFD
Females
% of field
p>0.99
p=0.10
p=0.86
M
HFD
Average lipid droplet size (µm²)
p=0.43
N
HFD
Lipid droplet count per 10x field
p=0.18
O
CHOW
Serum AST (U/L)
p=0.040
p=0.011
P
CHOW
Serum ALT (U/L)
p=0.0029
p=0.068
Q
HFD
Serum AST (U/L)
p=0.0015
p=0.030
R
HFD
Serum ALT (U/L)
p=0.044
p=0.23
S
T
Chow
HFD
100 µm
U
CHOW
Inflammation score (0-3)
p=0.59
p=0.54
V
HFD
Inflammation score (0-3)
p=0.031
p=0.64
W
Nuclei clusters per field
p=0.25
p=0.031
Chow
HFD

**Figure S2. Loss of EFHD1 prevents hepatocyte injury. A, B.** Serum triglyceride levels, normal chow (A, WT: males n=10, females n=7; *Efhd1*$^{-/-}$: males n=6, females n=8) or 28 weeks of HFD (B, WT: males n=6, females n=4; *Efhd1*$^{-/-}$: males n=4, females n=4). **C, D.** MASH activity score in normal chow (C) or 28 week HFD animals (D). **E, J.** Hepatocyte ballooning score in normal chow (E) or 28 week HFD animals (J). **F,G,K,L.** Histological assessment of liver fat in normal chow (F,G) or 28 week HFD animals (K,L). **H, I, M, N.** Direct measurement of lipid droplet size (H, M) and count (I, N). For normal chow, WT: n=19; *Efhd1*$^{-/-}$: n=23. For HFD, WT: n=28; *Efhd1*$^{-/-}$: n=16. **O-R.** Serum AST (O, Q) and ALT (P, R) measurements. For normal chow, WT: males n=11, females n=8; *Efhd1*$^{-/-}$: males n=10, females n=14. For HFD, WT: males n=11, females n=9; *Efhd1*$^{-/-}$: males n=21, females n=15. **S.** Representative liver micrographs following hematoxylin and eosin staining. Arrows indicate leukocyte clusters. **T.** Representative liver micrographs following Masson's trichrome staining. **U, V.** Histological assessment of inflammation for normal chow (U) or HFD (V). **W.** Direct assessment of leukocyte clusters from liver micrographs of normal chow (WT n=19, *Efhd1*$^{-/-}$ n=23) or HFD (WT n=19, *Efhd1*$^{-/-}$ n=23). All clinical histological scoring in C-G, J-L, U-V was assessed in blinded fashion by liver pathologist (K.J.E.). For normal chow in C, E-G, U, WT: males n=8, females n=11; *Efhd1*$^{-/-}$: males n=10, females n=13. For HFD in D, J-L, V, WT: males n=19, females n=9; *Efhd1*$^{-/-}$: males n=9, females n=7. Bar: mean ± SEM.

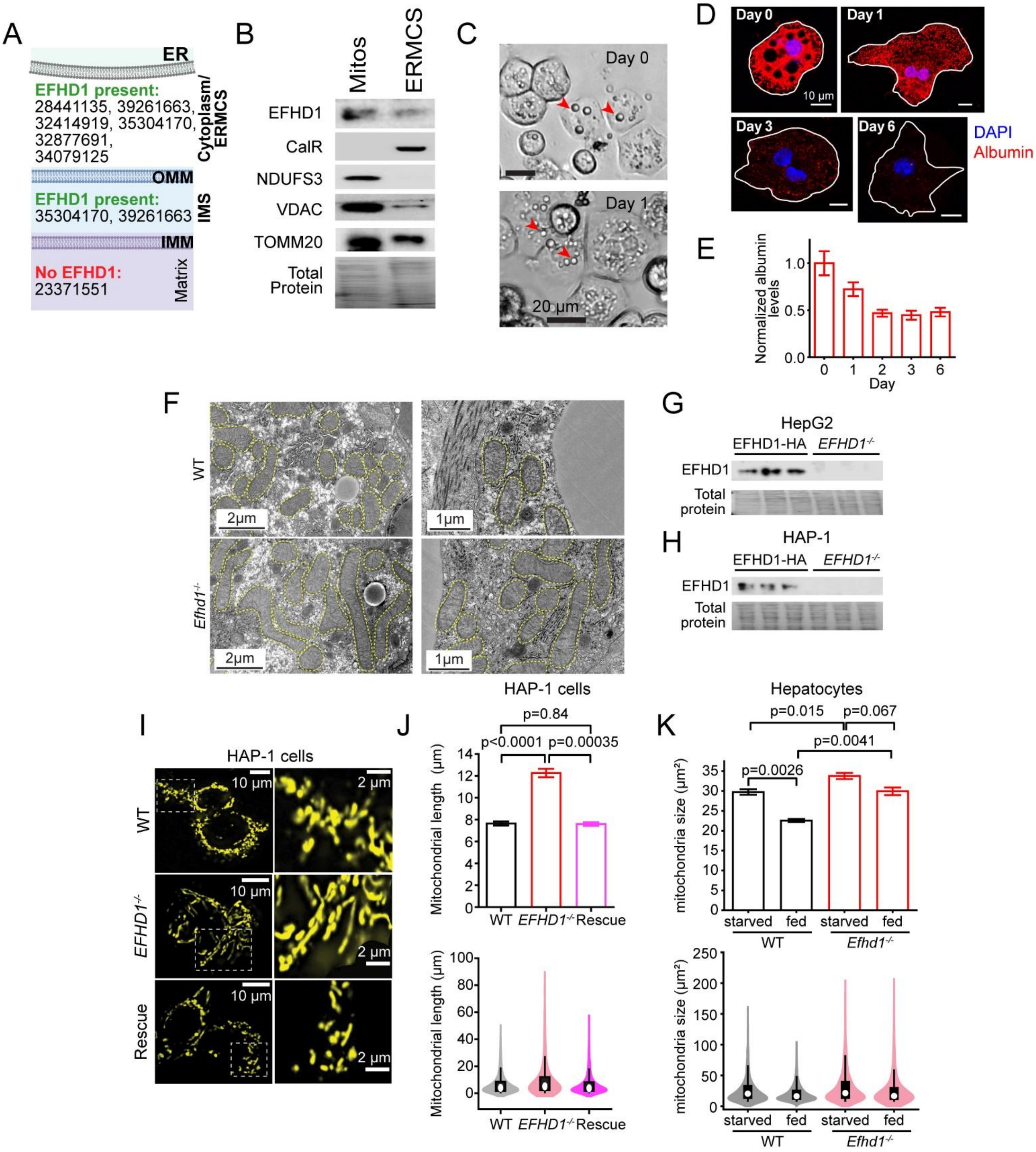

A
ER
EFHD1 present:
28441135, 39261663, 32414919, 35304170, 32877691, 34079125
Cytoplasm/ ERMCS
OMM
EFHD1 present:
35304170, 39261663
IMS
IMM
No EFHD1:
23371551
Matrix
B
Mitos
ERMCS
EFHD1
CalR
NDUFS3
VDAC
TOMM20
Total Protein
C
Day 0
Day 1
20 μm
D
Day 0
Day 1
Day 3
Day 6
10 μm
DAPI
Albumin
E
Normalized albumin levels
Day
F
WT
Efhd1-/-
2μm
1μm
G
HepG2
EFHD1-HA
EFHD1-/-
EFHD1
Total protein
H
HAP-1
EFHD1-HA
EFHD1-/-
EFHD1
Total protein
I
HAP-1 cells
WT
EFHD1-/-
Rescue
10 μm
2 μm
J
HAP-1 cells
p=0.84
p<0.0001
p=0.00035
Mitochondrial length (μm)
WT
EFHD1-/-
Rescue
Mitochondrial length (μm)
K
Hepatocytes
p=0.015
p=0.067
p=0.0041
p=0.0026
mitochondria size (μm²)
starved
fed
WT
Efhd1-/-
mitochondria size (μm²)

**Figure S3. Additional data for EFHD1 contribution to $Ca^{2+}$-induced mitochondrial remodeling. A.** Summary of proximity ligation and proteinase protection studies (listed by Pubmed ID), indicating that EFHD1 is primarily present in cytoplasm and IMS. EFHD1 is also present, though not necessarily enriched, at ERMCS. **B.** Mouse liver western blot analysis showing that EFHD1 is present in both mitochondrial and ERMCS fractions. CalR: Calreticulin (ER marker); NDUFS3: NADH:ubiquinone oxidoreductase core subunit S3 (mitochondrial matrix/IMM marker); VDAC: Voltage-dependent anion channel (OMM/ERMCS marker); TOMM20: translocase of outer mitochondrial membrane 20 (OMM/ERMCS marker).**C.** Representative bright field images of WT hepatocytes immediately after (day 0) and 24 hours (day 1) after isolation. Arrowheads indicate lipid droplets which differentiate hepatocytes from other liver cells. **D, E.** Representative images (D) and summary (E) showing albumin staining in hepatocytes on days 0, 1, 3 and 6 post isolation. Albumin levels decrease as hepatocytes de-differentiate. Day 0: n=22 hepatocytes from N=2 mice; Day 1: N=2, n=23; Day 2: N=2, n=21; Day 3: N=2, n=21; Day 6: N=2, n=24. All experimental assays were performed on isolated hepatocytes within 24 hours of isolation. **F.** Same image as Fig. 2D with mitochondria outlined to highlight changes in *Efhd1*$^{-/-}$ hepatocytes. **G, H.** Absent EFHD1 on western blot in *EFHD1*$^{-/-}$ HepG2 (G) or HAP-1 (H) cells compared with EFHD1 overexpressed cells (EFHD1-HA). We compared overexpressed (EFHD1-HA) to knockout (*EFHD1*$^{-/-}$) cells because of low endogenous EFHD1 expression in HepG2 cells. **I, J.** Representative images (I) and mitochondrial length quantification (J) of HAP-1 cells stained with 200 nM MitoTracker Orange. Right, boxed inset at higher magnification. Rescue is *EFHD1*$^{-/-}$ HAP-1 expressing EFHD1-HA. WT n=4821 mitochondria from N=99 cells; *EFHD1*$^{-/-}$ N=111, n=4368; EFHD1-HA N=104, n=4811. **K.** Comparison of mitochondrial size in WT or *Efhd1*$^{-/-}$ hepatocytes incubated for 24 hours in either low-glucose (1 g/L, starved) or high-glucose, high-fat media (4.5 g/L D-glucose, 0.4 mM palmitate, fed). WT fed n=3770 mitochondria from N=2 mice; WT starvation N=2 n=2530; *Efhd1*$^{-/-}$ fed N=2, n=6883; *Efhd1*$^{-/-}$ starvation N=2, n=4382. Data in J-K is displayed as a bar chart (top) to show mean effects and as a violin plot (bottom) to show the full distribution. Bars are mean ± SEM.

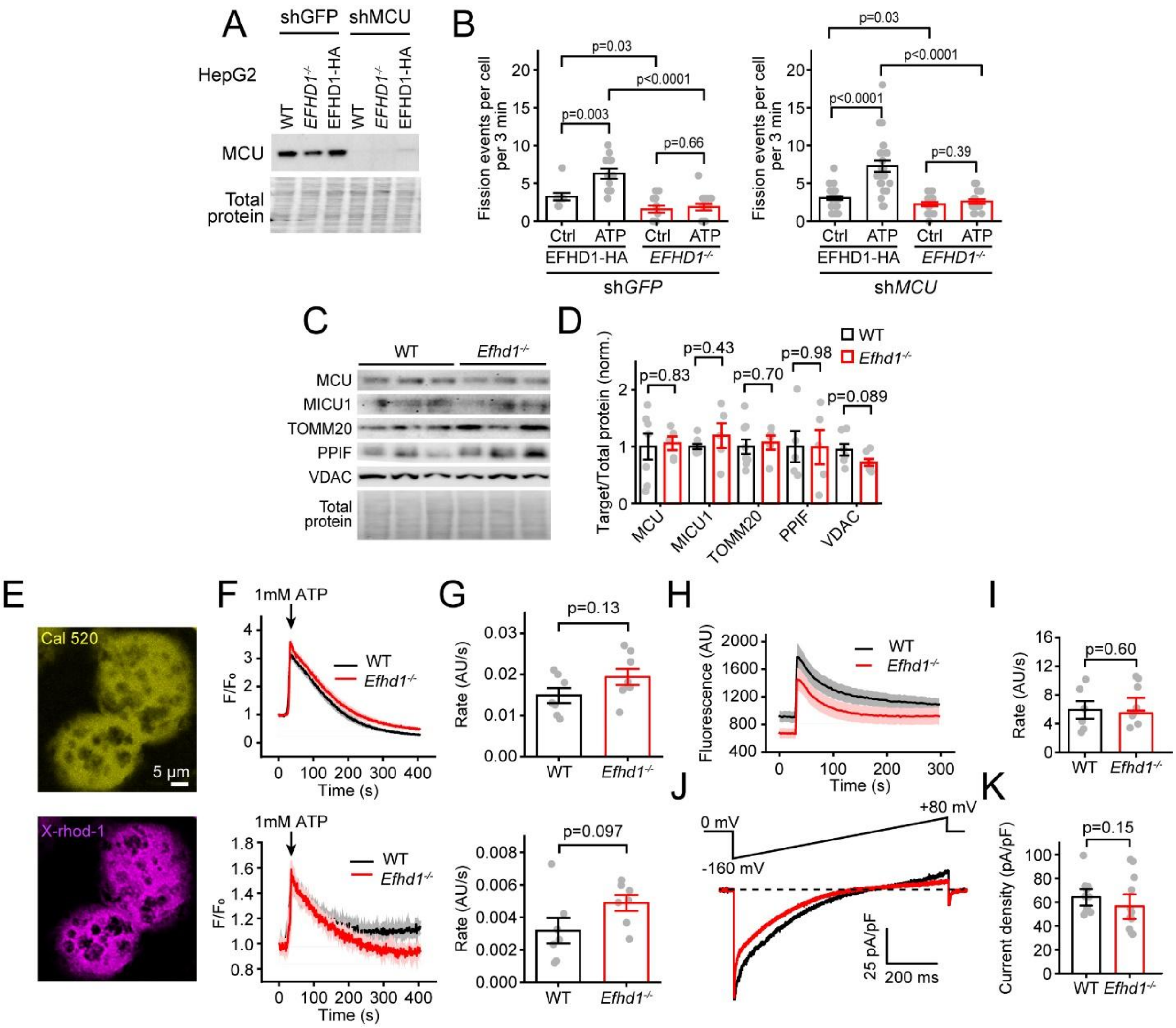


**Figure S4. EFHD1 effects on $Ca^{2+}$-dependent mitochondrial remodeling are independent of mitochondrial $Ca^{2+}$ uptake. A.** MCU levels are diminished in sh*MCU*-expressing HepG2 cells relative to control (sh*GFP*) on Western blot. **B.** Direct measurement of fission events after addition of vehicle (PBS, ctrl) or 1mM ATP in HepG2 cells as in Fig. 1L. Inhibiting MCU had no effect on the rate of ATP-induced fission events. shGFP: EFHD1-HA Ctrl: n=10; *Efhd1*$^{-/-}$ Ctrl: n=12; EFHD1-HA ATP: n=13; *Efhd1*$^{-/-}$ ATP: n=15. sh*MCU*: EFHD1-HA Ctrl: n=34; *Efhd1*$^{-/-}$ Ctrl: n=25; EFHD1-HA ATP: n=20; *Efhd1*$^{-/-}$ ATP: n=20. **C, D.** Western blot (C) and summary (D) of liver mitochondrial $Ca^{2+}$ homeostasis proteins MCU: mitochondrial $Ca^{2+}$ uniporter; MICU1: mitochondrial calcium uptake 1; TOMM20: translocase of outer mitochondrial membrane 20; PPIF: peptidylprolyl isomerase F; VDAC: voltage-dependent anion channel. TOMM20 is same as in Fig. 2B. MCU: N=7 WT and N=5 *Efhd1*$^{-/-}$ mice; MICU1: N=9 WT and N=5 *Efhd1*$^{-/-}$ mice; TOMM20: N=7 WT and N=5 *Efhd1*$^{-/-}$ mice; PPIF: N=5 WT and N=5 *Efhd1*$^{-/-}$ mice; VDAC: N=7 WT and N=7 *Efhd1*$^{-/-}$ mice. **E-G.** ATP-induced $Ca^{2+}$ uptake in isolated hepatocytes. **E.** Isolated hepatocytes stained with cytoplasmic $Ca^{2+}$ sensor Cal-520 (yellow, top) and mitochondrial $Ca^{2+}$

sensor X-rhod-1 (magenta, bottom). **F.** Cytoplasmic (top, Cal-520) and mitochondrial (bottom, X-rhod-1) $Ca^{2+}$ transients from isolated hepatocytes following treatment with 1 mM ATP. **G.** $Ca^{2+}$ uptake rate summary calculated from traces as in (d) (WT n=7; *Efhd1*$^{-/-}$ n=8). **H, I.** $Ca^{2+}$ uptake in purified liver mitochondria, which disrupts differences in mitochondrial networks between *Efhd1*$^{-/-}$ and WT. Traces (H) and summary (I) of $Ca^{2+}$ uptake measured via a decline in extramitochondrial $Ca^{2+}$ (1 μM Oregon Green BAPTA-6F) following a 10 μM $Ca^{2+}$ bolus (WT n=6, *Efhd1*$^{-/-}$ n=8). **J, K.** Whole-mitoplast electrophysiology to directly measure mitochondrial $Ca^{2+}$ currents. (J) Top, voltage ramp protocol. Bottom, exemplar $Ca^{2+}$ currents from isolated liver mitoplasts. (K) Summary. Assays in this figure were performed on normal chow-fed mice. Bars: mean ± SEM.

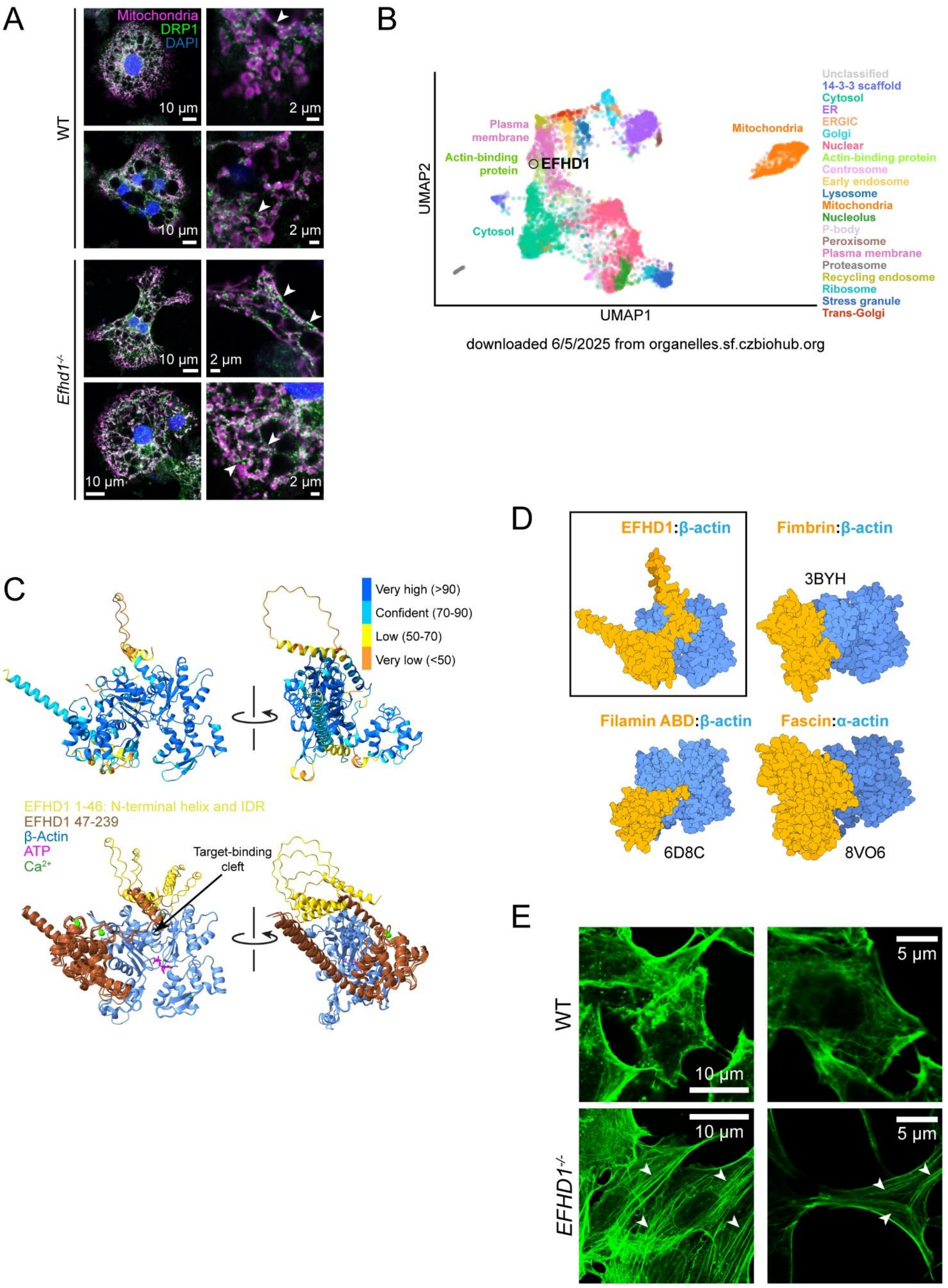
A
Mitochondria
DRP1
DAPI
WT
Efhd1-/-
10 µm
2 µm
B
UMAP2
UMAP1
Plasma membrane
Actin-binding protein
EFHD1
Cytosol
Mitochondria
Unclassified
14-3-3 scaffold
Cytosol
ER
ERGIC
Golgi
Nuclear
Actin-binding protein
Centrosome
Early endosome
Lysosome
Mitochondria
Nucleolus
P-body
Peroxisome
Plasma membrane
Proteasome
Recycling endosome
Ribosome
Stress granule
Trans-Golgi
downloaded 6/5/2025 from organelles.sf.czbiohub.org
C
Very high (>90)
Confident (70-90)
Low (50-70)
Very low (<50)
EFHD1 1-46: N-terminal helix and IDR
EFHD1 47-239
β-Actin
ATP
Ca2+
Target-binding cleft
D
EFHD1:β-actin
Fimbrin:β-actin
3BYH
Filamin ABD:β-actin
Fascin:α-actin
6D8C
8VO6
E
WT
EFHD1-/-
10 µm
5 µm

**Figure S5. EFHD1 promotes actin rearrangements. A.** Representative hepatocytes from normal chow-fed mice stained for mitochondria (magenta) and DRP1 (green). Arrowheads show mitochondria-localized DRP1 puncta. **B.** Native organelle immunoprecipitation data from PMID 39742809 annotates EFHD1 primarily as an actin-binding protein. **C.** Top, Ribbon diagram depiction of AlphaFold3 confidence (predicted local distance difference test, pLDDT) for EFHD1:β-actin structure. pLDDT at the interface between EFHD1 and β-actin is >90. Bottom, in the overlay of the top 5 AlphaFold3 predicted structures, EFHD1 bound β-actin via a helical sequence linking the IDR to the EF hands, with some further interactions extending to the proximal portion of the coiled-coil domain. The greatest uncertainty is in the localization of the N-terminal helix and IDR, with some structures showing it occupying the target-binding cleft in the actin + end where polymerization occurs. **D.** Comparison of the AlphaFold3 predicted EFHD1:β-actin structure with other actin-bundling proteins shows all bind at a similar location at actin domain 1-2. PDB structure IDs are shown for fimbrin, filamin actin-binding domain (ABD) and fascin. **E.** Representative images showing actin (green) morphology via phalloidin staining in HAP-1 cells. Arrowheads indicate prominent cytoplasmic actin stress fibers observed in *EFHD1*$^{-/-}$ cells.

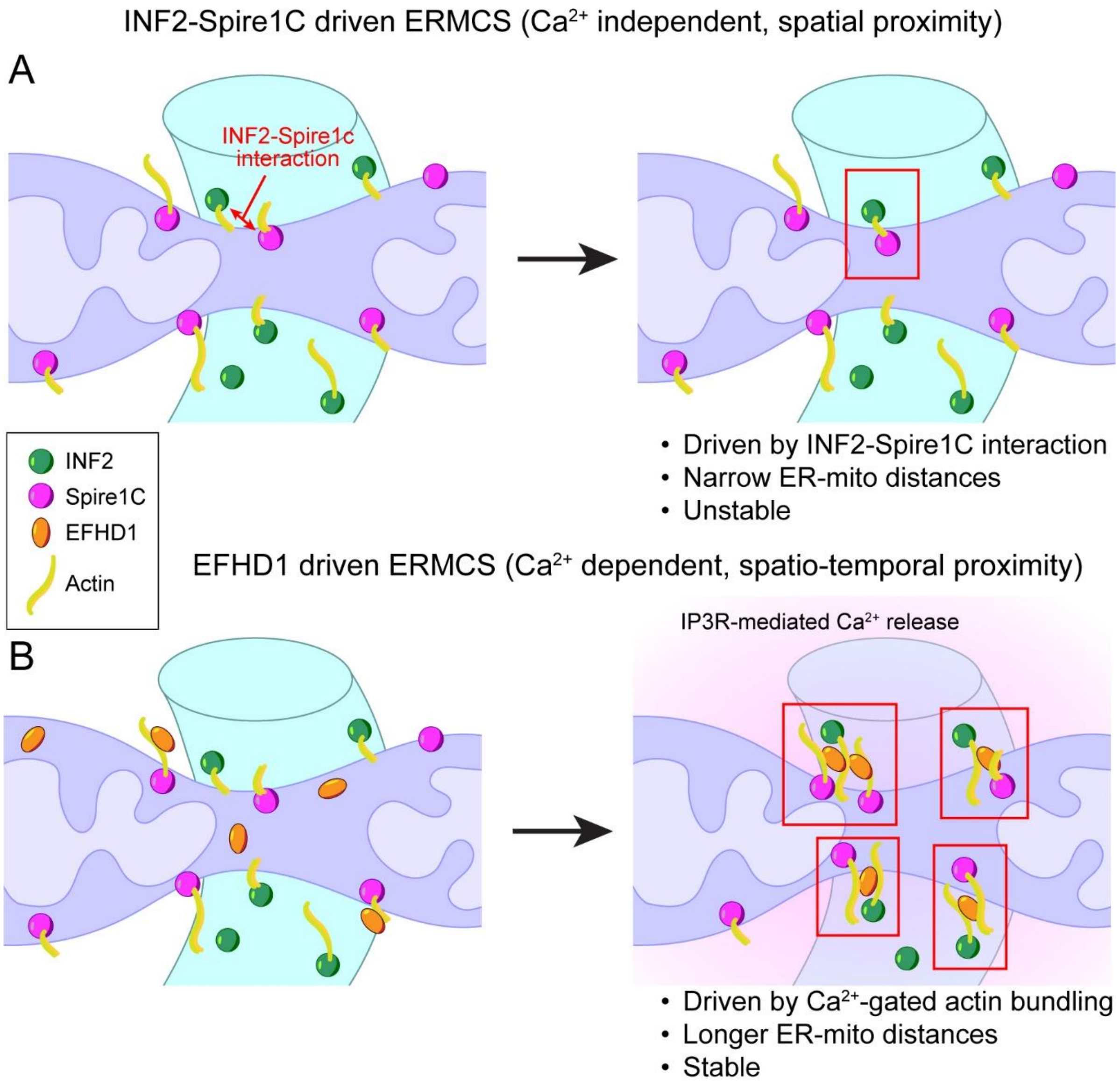


**Figure S6. Working model for EFHD1-driven $Ca^{2+}$-dependent ERMCS stabilization. A.** The current model for actin crosslinking at ERMCS requires a direct interaction between the OMM-bound actin-nucleating factor Spire1C, and the ER-bound actin polymerization factor INF2. Interactions between these two factors are possibly less efficient and unstable as they are localized on opposite membranes and may have limited contact. In the absence of EFHD1, ERMCS are narrower because of the requirement for more direct interactions. **B.** EFHD1-driven fission does not require direct interaction between ER- and OMM-bound factors. Instead, actin filaments may be more diffuse and motile at ERMCS. Upon $Ca^{2+}$ release from the ER, EFHD1 is poised to integrate spatiotemporal proximity to crosslink actin filaments originating from opposite membranes, leading to more stable actin-driven constriction at ERMCS.

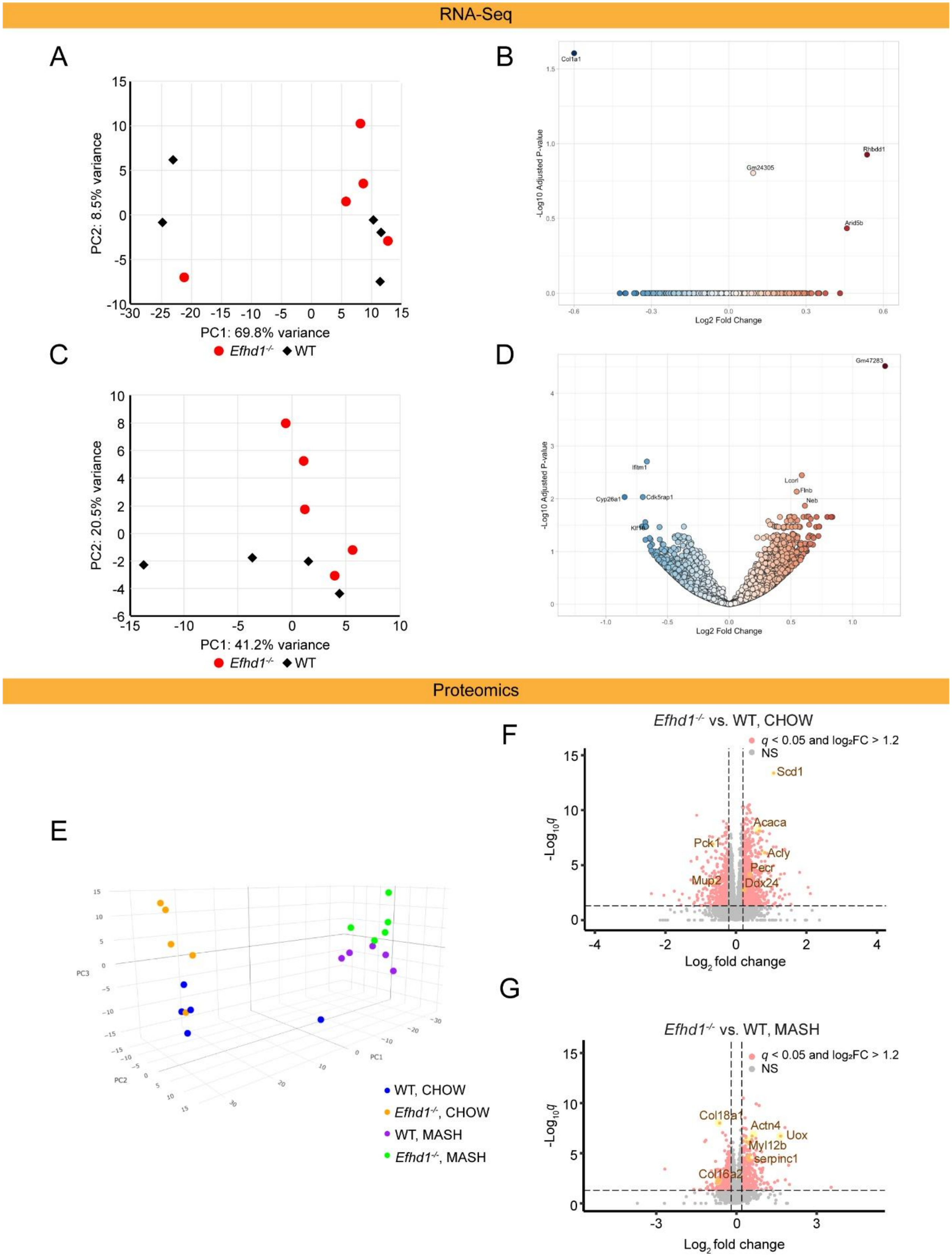
RNA-Seq
A
PC2: 8.5% variance
PC1: 69.8% variance
Efhd1-/- WT
B
Col1a1
Rhbdd1
Gm24305
Arid5b
-Log10 Adjusted P-value
Log2 Fold Change
C
PC2: 20.5% variance
PC1: 41.2% variance
Efhd1-/- WT
D
Gm47283
Ifitm1
Cyp26a1
Cdk5rap1
Lcorl
Flnb
Neb
-Log10 Adjusted P-value
Log2 Fold Change
Proteomics
E
PC1
PC2
PC3
WT, CHOW
Efhd1-/-, CHOW
WT, MASH
Efhd1-/-, MASH
F
Efhd1-/- vs. WT, CHOW
q < 0.05 and log2FC > 1.2
NS
Scd1
Acaca
Pck1
Acly
Pecr
Mup2
Ddx24
-Log10q
Log2 fold change
G
Efhd1-/- vs. WT, MASH
q < 0.05 and log2FC > 1.2
NS
Col18a1
Actn4
Uox
Myl12b
serpinc1
Col16a2
-Log10q
Log2 fold change

**Figure S7. Further transcriptomic and proteomic analysis of EFHD1 ablation. A,B.** From whole liver RNA-seq of *Efhd1*$^{-/-}$ versus WT mice fed normal chow, (A) principal components analysis using the top 500 most variable genes, and (B) volcano plot of differential gene expression. **C,D.** From whole liver RNA-seq of *Efhd1*$^{-/-}$ versus WT mice fed a MASH diet, (A) principal components analysis using the top 500 most variable genes, and (B) volcano plot of differential gene expression. **E-G.** From whole liver proteomics of *Efhd1*$^{-/-}$ versus WT mice fed a normal chow or MASH diet, (E) principal components analysis using proteins expressed in all samples, and volcano plot of differential gene expression comparing *Efhd1*$^{-/-}$ versus WT mice fed a (F) normal chow or (G) MASH diet. RNA-seq, chow: WT n=5, *Efhd1*$^{-/-}$ n=5. MASH: WT n=4, *Efhd1*$^{-/-}$ n=5. Proteomics, chow: WT n=5, *Efhd1*$^{-/-}$ n=5. MASH: WT n=5, *Efhd1*$^{-/-}$ n=5.

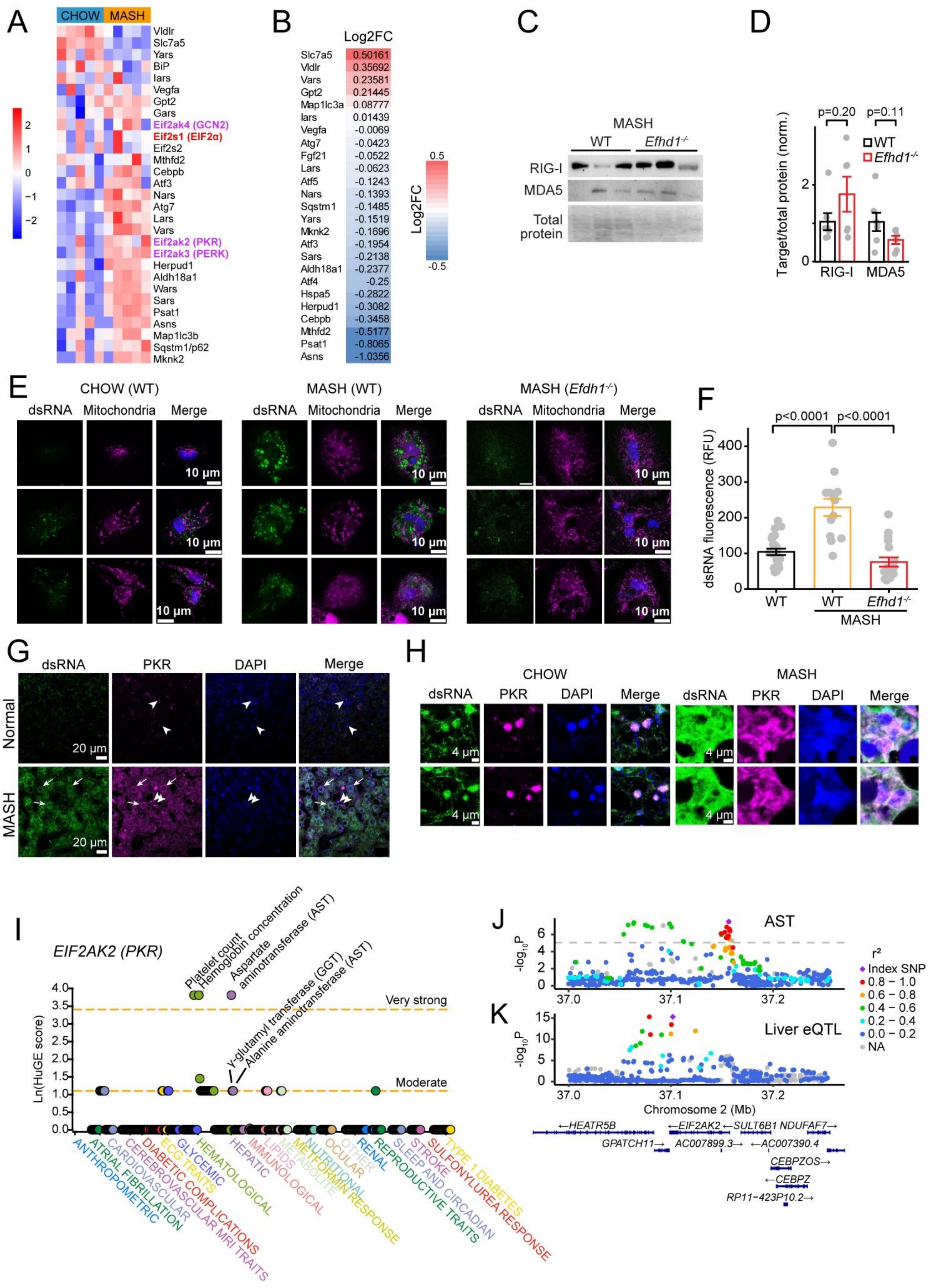

A
CHOW
MASH
Vldlr
Slc7a5
Yars
BiP
Iars
Vegfa
Gpt2
Gars
Eif2ak4 (GCN2)
Eif2s1 (EIF2α)
Eif2s2
Mthfd2
Cebpb
Atf3
Nars
Atg7
Lars
Vars
Eif2ak2 (PKR)
Eif2ak3 (PERK)
Herpud1
Aldh18a1
Wars
Sars
Psat1
Asns
Map1lc3b
Sqstm1/p62
Mknk2
B
Log2FC
Slc7a5 0.50161
Vldlr 0.35692
Vars 0.23581
Gpt2 0.21445
Map1lc3a 0.08777
Iars 0.01439
Vegfa -0.0069
Atg7 -0.0423
Fgf21 -0.0522
Lars -0.0623
Atf5 -0.1243
Nars -0.1393
Sqstm1 -0.1485
Yars -0.1519
Mknk2 -0.1696
Atf3 -0.1954
Sars -0.2138
Aldh18a1 -0.2377
Atf4 -0.25
Hspa5 -0.2822
Herpud1 -0.3082
Cebpb -0.3458
Mthfd2 -0.5177
Psat1 -0.8065
Asns -1.0356
C
MASH
WT
Efhd1-/-
RIG-I
MDA5
Total protein
D
p=0.20
p=0.11
WT
Efhd1-/-
Target/total protein (norm.)
RIG-I
MDA5
E
CHOW (WT)
MASH (WT)
MASH (Efdh1-/-)
dsRNA
Mitochondria
Merge
10 μm
F
p<0.0001
p<0.0001
dsRNA fluorescence (RFU)
WT
WT
Efhd1-/-
MASH
G
dsRNA
PKR
DAPI
Merge
Normal
MASH
20 μm
H
CHOW
MASH
dsRNA
PKR
DAPI
Merge
4 μm
I
EIF2AK2 (PKR)
Platelet count
Hemoglobin concentration
Aspartate aminotransferase (AST)
γ-glutamyl transferase (GGT)
Alanine aminotransferase (AST)
Very strong
Moderate
Ln(HuGE score)
ANTHROPOMETRIC
ATRIAL FIBRILLATION
CARDIOVASCULAR
CEREBROVASCULAR MRI TRAITS
DIABETIC COMPLICATIONS
ECG TRAITS
GLYCEMIC
HEMATOLOGICAL
HEPATIC
IMMUNOLOGICAL
LIPIDS
METABOLITE
METFORMIN RESPONSE
NUTRITIONAL
OCULAR
OTHER
RENAL
REPRODUCTIVE TRAITS
SLEEP AND CIRCADIAN
STROKE
SULFONYLUREA RESPONSE
TYPE 1 DIABETES
J
AST
-log10P
r²
Index SNP
0.8 − 1.0
0.6 − 0.8
0.4 − 0.6
0.2 − 0.4
0.0 − 0.2
NA
K
Liver eQTL
Chromosome 2 (Mb)
←HEATR5B
←EIF2AK2
←SULT6B1
NDUFAF7→
GPATCH11→
AC007899.3→
←AC007390.4
CEBPZOS→
←CEBPZ
RP11-423P10.2→

**Figure S8. Pathways to the integrated stress response in MASH. A.** Heat maps of protein expression of ATF4 target genes from livers of WT mice fed normal chow or MASH diet. **B.** Heat map of ATF4 target gene expression from RNA-seq data of *Efhd1*$^{-/-}$ versus WT mice fed MASH diet. Most targets are downregulated (negative) in *Efhd1*$^{-/-}$ relative to WT (WT n=4, *Efhd1*$^{-/-}$ n=5). **C, D.** Western blot (C) and summary (D) of RIG-I and MDA5 liver expression. RIG-I: Retinoic acid-inducible gene I; MDA5: Melanoma Differentiation-Associated gene 5 (WT n=5, *Efhd1*$^{-/-}$ n=5). **E.** Additional representative images, as in Fig. 3G, of dsRNA and mitochondrial staining in isolated mouse hepatocytes. **F.** Quantification of dsRNA fluorescence (WT chow n=23 from 4 mice, WT MASH n=13 from 5 mice, *Efhd1*$^{-/-}$ MASH n=18 from 5 mice). **G.** Additional exemplars as in Fig. 3H. Arrowheads show PKR staining in dense non-hepatocyte nuclei, while arrows show dsRNA- and PKR-stained cytoplasmic signal. **H.** Additional representative images as in Fig. 3I. **I.** The panel was downloaded and modified from https://hugeamp.org/gene.html?gene=EIF2AK2 on Feb 8, 2025, and represents the calculated Human Genetic Evidence (HuGE) score, which quantifies genetic support for involvement in the diseases and traits available in the Common Metabolic Diseases Knowledge Portal. Shown are different thresholds (dotted lines) for the quality of evidence. **J, K.** *EIF2AK2* (PKR) association with circulating aspartate aminotransferase (AST) (J) in UK Biobank and liver eQTL (K). Lead variant, purple diamond. Bars: mean ± SEM.

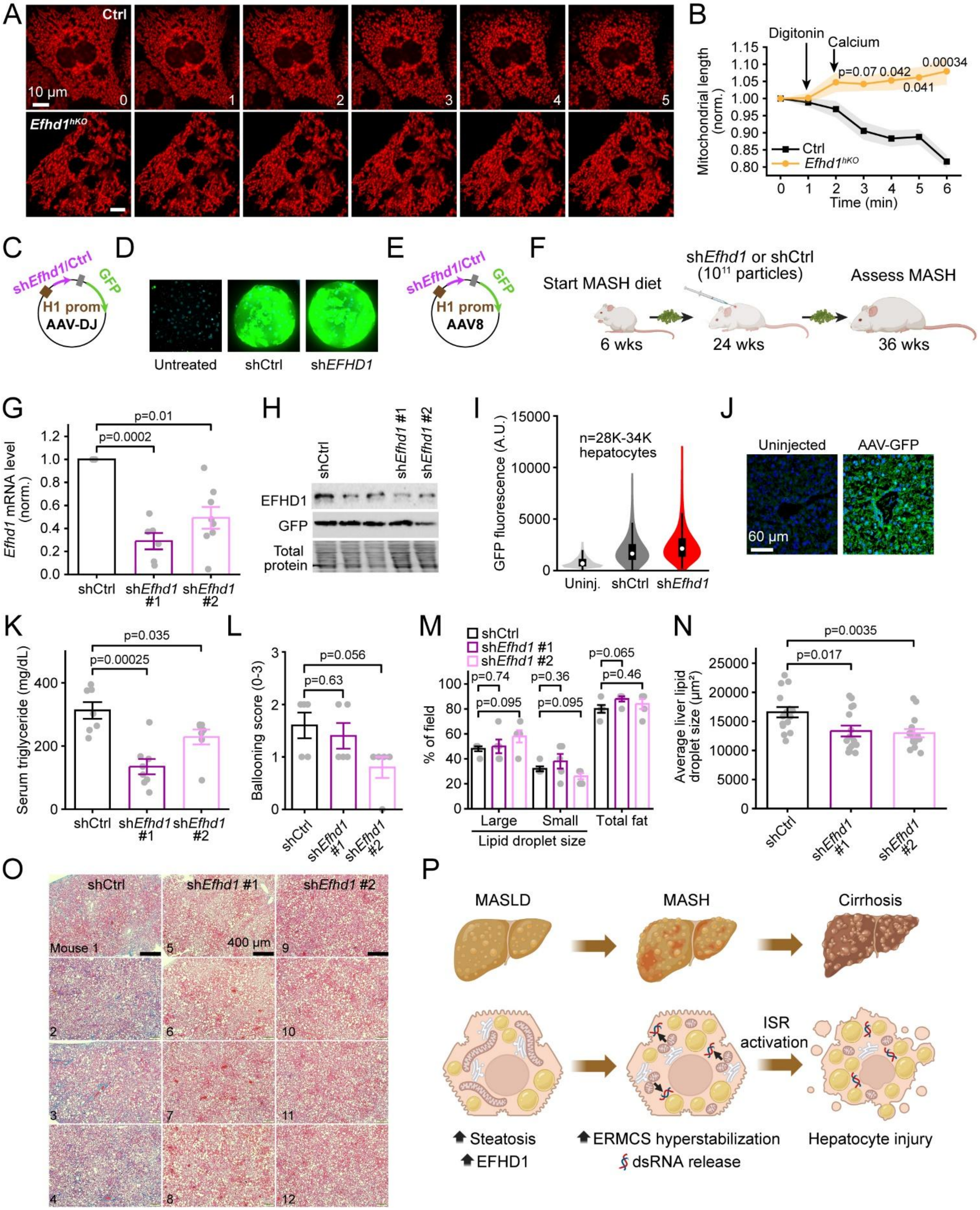

A
Ctrl
10 μm
0
1
2
3
4
5
Efhd1^hKO
B
Digitonin
Calcium
p=0.07 0.042
0.00034
0.041
Mitochondrial length (norm.)
Ctrl
Efhd1^hKO
Time (min)
C
shEfhd1/Ctrl
GFP
H1 prom
AAV-DJ
D
Untreated
shCtrl
shEFHD1
E
shEfhd1/Ctrl
GFP
H1 prom
AAV8
F
Start MASH diet
shEfhd1 or shCtrl (10^11 particles)
Assess MASH
6 wks
24 wks
36 wks
G
p=0.01
p=0.0002
Efhd1 mRNA level (norm.)
shCtrl
shEfhd1 #1
shEfhd1 #2
H
shCtrl
shEfhd1 #1
shEfhd1 #2
EFHD1
GFP
Total protein
I
n=28K-34K hepatocytes
GFP fluorescence (A.U.)
Uninj.
shCtrl
shEfhd1
J
Uninjected
AAV-GFP
60 μm
K
p=0.035
p=0.00025
Serum triglyceride (mg/dL)
shCtrl
shEfhd1 #1
shEfhd1 #2
L
p=0.056
p=0.63
Ballooning score (0-3)
shCtrl
shEfhd1 #1
shEfhd1 #2
M
shCtrl
shEfhd1 #1
shEfhd1 #2
p=0.74
p=0.095
p=0.36
p=0.095
p=0.065
p=0.46
% of field
Large
Small
Total fat
Lipid droplet size
N
p=0.0035
p=0.017
Average liver lipid droplet size (μm²)
shCtrl
shEfhd1 #1
shEfhd1 #2
O
shCtrl
shEfhd1 #1
shEfhd1 #2
Mouse 1
400 μm
P
MASLD
MASH
Cirrhosis
ISR activation
Steatosis
EFHD1
ERMCS hyperstabilization
dsRNA release
Hepatocyte injury

**Figure S9. Liver-specific EFHD1 inhibition. A, B**. Representative images (A) and summary of mitochondrial length (B) in MitoTracker-labeled isolated mouse hepatocytes. Cells were imaged for 6 minutes, and treated sequentially with digitonin and a $Ca^{2+}$ bolus. **C.** Schematic of AAV plasmid used for acute EFHD1 inhibition in human liver organoids. **D.** Maximum intensity projection image of GFP expression showing successful organoid AAV transduction. **E.** Schematic of AAV plasmid used for acute EFHD1 inhibition in mouse liver. **F.** Timeline of AAV8-mediated *Efhd1* inhibition experiments. Image created using BioRender. **G.** Quantitative RT-PCR of liver *Efhd1* mRNA level after AAV8 treatments. **H.** Confirmation of AAV8-mediated EFHD1 inhibition via Western blot of livers. Middle two unlabeled lanes are from other sh*Efhd1* constructs screened but not used further. **I.** GFP fluorescence measured via flow cytometry from hepatocytes isolated from uninjected or AAV8-treated mice. **J.** Representative images comparing GFP fluorescence from liver sections of uninjected mice and mice injected with AAV8. **K.** Serum triglyceride levels. **L.** Hepatocyte ballooning score. **M.** Histopathological assessment of liver fat. **N.** Direct measurement of lipid droplet size. **O.** Additional representative liver micrographs following Masson's trichrome staining, with fibrosis evident as blue stain. Each panel is from a different injected mouse. **P.** Proposed mechanism for EFHD1-dependent pathological mitochondrial remodeling in MASH. Created with BioRender. For G, shCtrl n=3; sh*Efhd1#1* n=6; sh*Efhd1#2* n=8. For K, shCtrl n=7; sh*Efhd1#1* n=8; sh*Efhd1#2* n=7. For L, M, shCtrl n=5; sh*Efhd1#1* n=5; sh*Efhd1#2* n=5. For N, shCtrl n=15; sh*Efhd1#1* n=15; sh*Efhd1#2* n=15. Bar: mean ± SEM.

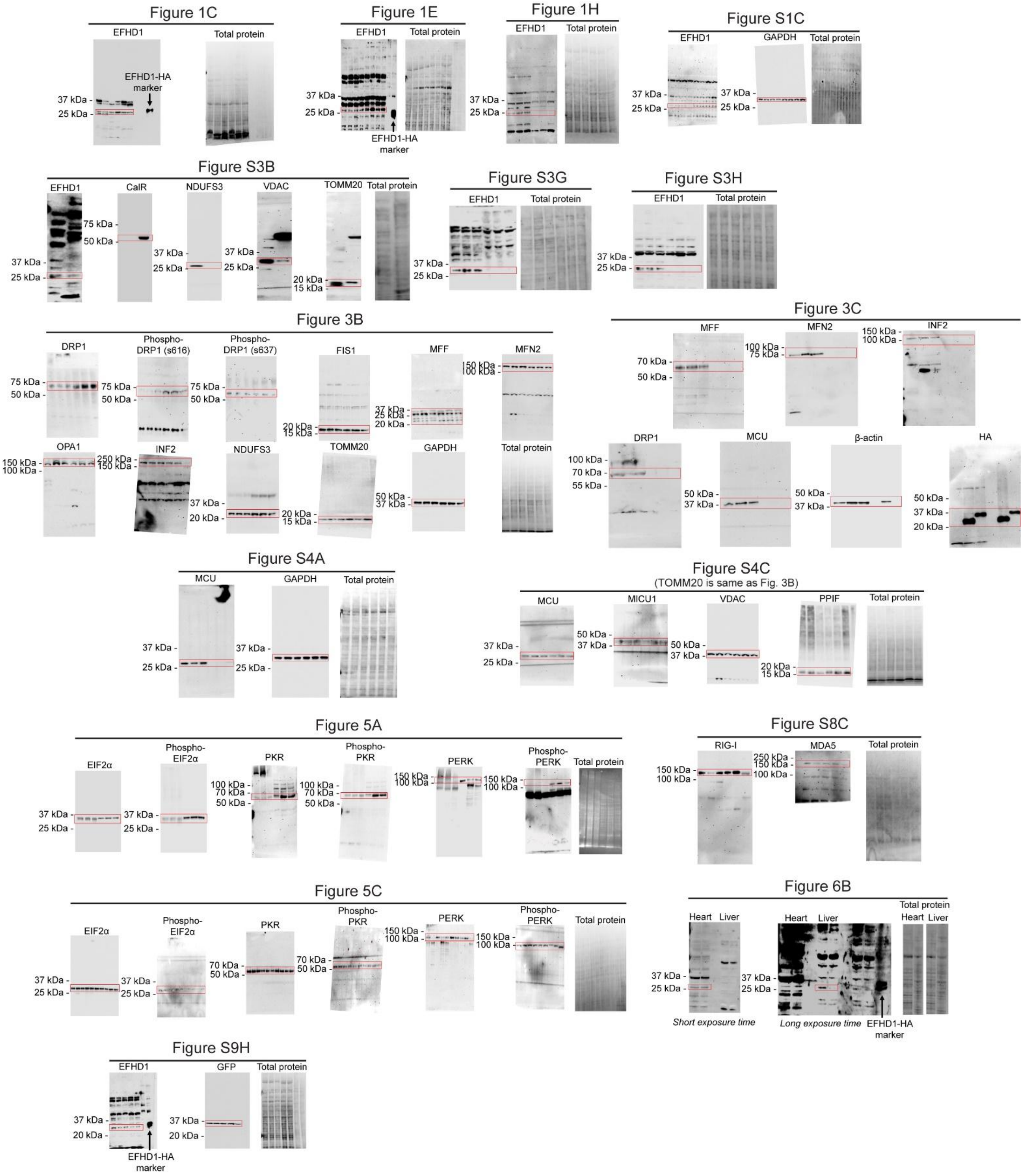


**Figure S10.** Uncropped images of all Western blots.

### SUPPLEMENTAL METHODS

#### Study approval

The study protocol for human liver tissue sections was reviewed by the Institutional Review Board at HCI and determined to be exempt from human subjects research (exemption number IRB_00091019). All animal procedures have been reviewed and approved by the Institutional Animal Care and Use Committee at the University of Utah, and for the whole-mitoplast electrophysiology experiments, at the University of Maryland Baltimore.

#### Antibodies

Albumin (Bethyl Laboratories,A90-234A,1:200 dilution immunofluorescence [IF]), β-actin (CST, 4970L, 1:1000 dilution Western blot [WB]), Calreticulin (CST,12238T, 1:1000 dilution WB), DRP1 (CST, D6C7, 1:1000 dilution WB, 1:200 dilution IF), DRP1 (Proteintech, 12957-1-AP, 1:1000 dilution WB), pDRP1 s637 (CST, 4867S, 1:1000 dilution WB), dsRNA (Sigma, MABE1134, 1:200 dilution IF), EFHD1 (Novus, NBP2-33281, 1:200 dilution IF), EIF2a (CST, 5324S, 1:1000 dilution WB), FIS1 (Proteintech,10956-1-AP, 1:1000 dilution WB, 1:200 dilution IF), GAPDH (CST, 2118L, 1:1000 dilution WB), GFP (Addgene, 180084-rAb, 1:1000 dilution WB), Goat anti-mouse IgG Alexa 488 (Thermo , A32723, 1:10000 dilution WB), Goat anti-rabbit IgG Alexa 555 (Thermo, A32740, 1:10000 dilution WB), HA (CST, 3724S, 1:1000 dilution WB), INF2 (Proteintech, 20466-1-AP, 1:1000 dilution WB, 1:200 dilution ICC), MCU (CST, 14997S, 1:1000 dilution WB), MDA5 (Proteintech, 21775-1-AP, 1:1000 dilution WB), MFF (CST, 84580S, 1:1000 dilution WB), MFN2 (CST, 11925T, 1:1000 dilution WB), NDUFS3 (Abcam, ab110249, 1:1000 dilution WB), OPA1 (CST, 80471T, 1:1000 dilution WB), PDH (CST, 3205T, 1:1000 dilution WB), p-PDH (Abcam, ab177461, 1:1000 dilution WB), pDRP1 s616 (CST, 3455S, 1:1000 dilution WB), p-EIF2a (CST, 3398T, 1:1000 dilution WB), PERK (CST, 3192S, 1:1000 dilution WB), p-PERK (Bioss Antibodies, bs-23340R-HRP, 1:1000 dilution WB), PKR (CST, 2297S, 1:1000 dilution WB), PKR (Proteintech, 18244-1-AP, 1:1000 dilution WB), PKR (Santa Cruz, sc-6282, 1:200 dilution ICC), p-PKR (Thermo, 44-668G, 1:1000 dilution WB), PPIF (Abcam, ab110324, 1:1000 dilution WB), RIG-I (CST, 3743S, 1:1000 dilution WB), RIG-I (Proteintech, 20566-1-AP, 1:1000 dilution WB), TOMM20 (CST, 42406T, 1:1000 dilution WB), VDAC (CST, 4661T, 1:1000 dilution WB). The EFHD1 rabbit polyclonal antibody was developed by Pacific Immunology, (Ramona, CA), and validated against *Efhd1*$^{-/-}$ lysates as described before(1). Rabbits were immunized with the EFHD1 peptide sequence: Cys-EQEERKREEEARRLRQAAFRELKAAFSA (Mouse *Efhd1* 212–240). The EFHD1 polyclonal antibody was used at a concentration of 1:500 for Western blot analysis.

#### Human liver samples

Liver tissue sections were from archival samples from deidentified patients seen at Huntsman Cancer Institute (Salt Lake City, UT). All patients were classified to their respective diagnosis by a pathologist at the time of initial collection. The diagnosis for individual tissue samples was confirmed by a pathologist based on histology review of formalin-fixed, paraffin-embedded sections taken from the same location as the tissue analyzed. The study protocol was reviewed by the Institutional Review Board at HCI and determined to be exempt from human subjects research (exemption number IRB_00091019).

#### *EFHD1*$^{-/-}$ cells

*EFHD1*$^{-/-}$ HepG2 and HAP-1 cells (Horizon Discovery) were generated using CRISPR Cas9. CRISPR components were obtained from Thermo (TrueCut Cas9 Protein v2 [A36499] and EFHD1 gRNA sequences: CCTGCAGGTATGACGCTGGG and TCGATGGCAAGCTCAGCTTC). CRISPR components were transfected into cells using electroporation (Neon System, Thermo Fisher) per the manufacturer's instructions. Individual clones were isolated by limiting dilution and knockout was confirmed via PCR and western blot before propagating.

**Gene Expression Omnibus Dataset analysis**

We downloaded cell summary data as indicated below and summed how many hepatocytes or non-hepatocytes expressed EFHD1.

| Dataset | Location: Filename Analyzed |
|---|---|
| GSE115469 | NCBI GEO: GSE115469_Data.csv and GSE115469_CellClusterType.txt |
| GSE124395 | NCBI GEO: GSE124395_Normalhumanlivercellatlasdata.txt and GSE124395_clusterpartition.txt supplementary files |
| GSE185477 | https://cellxgene.cziscience.com/collections/44531dd9-1388-4416-a117-af0a99de2294 : Healthy human liver: integrated - 10x 3 |
| GSE192740 | https://livercellatlas.org/download.php : gene counts and cell annotations for Human All Liver cell |
| GSE243977 | https://cellxgene.cziscience.com/collections/0c8a364b-97b5-4cc8-a593-23c38c6f0ac5 : All cells types from snRNA-seq of human primary sclerosing cholangitis patients and healthy controls" for both 10x 3' and 10x 5' |

**3D human liver microtissues**

Experiments were performed at InSphero (Switzerland) using 3D InSight Human Liver Microtissues as described previously(2). Microtissues were generated by self-assembly of a pool of monodispersed primary human hepatocytes from 10 donors, single-donor hepatic stellate cells and single-donor Kupffer and liver endothelial cells. After aggregation, microtissues were cultured in basal hepatocyte maintenance medium (CS-07-302-01, InSphero AG), with media exchange every 2-3 days. TNFα in microtissue supernatants was determined on day 5 using the Magnetic Luminex Performance Assay according to the manufacturer's instructions with adjustment of microparticle and antibody concentrations to analyte levels present in supernatants. Intracellular triglyceride content was measured with the Triglyceride-Glo assay (Promega), using a Tecan Spark 10 M plate reader.

**Mouse strains and animal handling**

All animal procedures have been reviewed and approved by the Institutional Animal Care and Use Committee at the University of Utah, and for the whole-mitoplast electrophysiology experiments, at the University of Maryland Baltimore. *Efhd1*$^{-/-}$ mice were obtained from the Jackson Laboratory (Bar Harbor, ME, C57BL/6NJ-Efhd1em1(IMPC)J/Mmjax, strain #042149)(1). Animals were kept on a C57BL/6NJ background, and wild-type mice of this strain were used as controls. *Efhd1*$^{hKO}$ mice were generated using the Cre-loxP system. The original strain contained a conditional-ready, lacZ-tagged mutant allele (JN958097.1) obtained from Helmholtz Zentrum München (IKMC Project 30211, ES cell clone EPD0370_1_C05, strain C57BL/6N-$^{Atm1Brd}$/a). The lacZ and neo cassettes were removed by crossing with a FLPo deleter strain (Jackson Laboratory strain #012930)(3), creating a conditional-ready floxed allele around exon 2. Cre-mediated deletion of exon 2 leads to removal of part of the EF hands as well as a frameshift that leads to early termination. The *Alb*-Cre mouse were obtained from Jackson Laboratory (strain # 003574)(4). Animals were backcrossed into the C57BL/6NJ strain. *Alb*-Cre mice were used as controls, to confirm that effects were not due to exogenous Cre expression. For AAV8 studies, we obtained C57BL/6NTac male mice fed a Gubra-Amylin MASH diet from 6 weeks of age

(MASH-B6-M, Taconic Biosciences, Germantown, NY). The mice were housed by Taconic until they were 20 weeks of age then transferred to the University of Utah CMC. They continued the MASH diet and were allowed to adapt for another 4 weeks at the University of Utah vivarium before proceeding with viral injections.

Animals were housed under standard conditions and allowed free access to food and water unless the experiment required fasting. Fasting was for no longer than 4 hours prior to euthanasia. Mice were euthanized via $CO_2$ inhalation with a second method performed such as cervical dislocation or removal of the heart of liver.

**MASH and high fat diets**

Mice were weaned at 3 week and fed a normal chow diet containing 10% fat until they were 8 weeks of age. At 8 weeks mice were transitioned from a normal chow diet to either a high fat diet or a MASH diet at 8 weeks of age. The high fat diet contained 60% fat (w/w, Envigo Adjusted Calories Diet (60/Fat), TD06414) and the MASH diet contained 40% Kcal Fat (Palm Oil), 2% Cholesterol, 20% Kcal/Fructose (Envigo, TD200591). Mice were fed high fat and MASH diets for 28 weeks and then sacked for experimental analysis.

**Carbon tetrachloride ($CCl_4$) injections**

Mice fed a normal chow diet were injected with 2 µL/g $CCl_4$ (Sigma, 56-23-5) suspended in corn oil (10%, v/v) via an Intraperitoneal (IP) injection 3 times per week for 6 weeks and then sacked for experimental analysis.

**Adeno-associated virus**

Self-complementary AAV particles were prepared by PackGene (Houston, TX) with low endotoxin titer (<10 EU/mL).

For human microtissues, shRNA sequences were packaged in AAV-DJ virus. Virus were applied to cells during microtissue aggregation at 3 x $10^5$ genome copies (GC) per microtissue.
Non-targeting: TCTCGCTTGGGCGAGAGTAAGtcaagagCTTACTCTCGCCCAAGCGAGA
sh*EFHD1*: GCAGAGTTGAAAGCTGAGCAAatattcaTTGCTCAGCTTTCAACTCTGC

Mice fed a MASH diet for 18 weeks (24 weeks of age) were injected with AAV8 virus containing shRNA targeted to either non-targeting control (sequence based on Sigma SHC216) or to two sequences targeting EFHD1. AAV8 virus was diluted 1:10 in PBS to a final concentration of $10^{12}$ genome copies/mL and 100 µL was injected into each mouse via tail vein injection. Mice were sacked 12 weeks later for analysis.
sh*Efhd1* #1: GTAAGTTCGAAGCTGAGTTAActcgagTTAACTCAGCTTCGAACTTAC,
sh*Efhd1* #2: GCTGGAAGGGACGGCTTTATTctcgagAATAAAGCCGTCCCTTCCAGC).

**Blood serum collection**

Mice were euthanized by $CO_2$ inhalation and blood samples collected via cardiac puncture by syringe with a 25G needle shortly after euthanasia. Blood was then spun at 2000 x *g* for 20 minutes and the serum collected. AST, ALT and triglyceride levels were measured using an Element DC chemistry analyzer (Antech Diagnostics) or via a Colorimetric Activity Assay Kit (Cayman Chemicals).

**Body composition analysis**

Mice were weighed and placed in a Bruker LF50 Time-Domain Nuclear Magnetic Resonance reader and their % body fat, % lean tissue and % fluid were measured and recorded.

**Lipid in stool**

Feces was collected from individual mice over 3–4 days and weighed. The feces was ground into a powder using a tissue grinder and 5 mL of normal saline (0.9% NaCl) was added to the powdered feces in a 15 mL tube and vortexed thoroughly. 5 mL of chloroform:methanol (2:1) was added to the tube and vortexed. The tube was centrifuged at 1000 x *g* for 10 min at room temperature. After centrifugation, two liquid phases were observed, the lower phase was removed using a 22G needle and transferred to a pre-weighed glass tube. The solvent was evaporated overnight in a fume hood and the dried lipid residue was weighed.

**Metabolic cages**

Mice aged 13 weeks fed either a normal chow diet or HFD for 5 weeks were placed in a metabolic chamber for 72 hours with a 12-hour light-dark cycle. $O_2$ consumption, $CO_2$ production, food and water intake and movement was monitored. These experiments were performed by the University of Utah Metabolic Phenotyping Core.

**Glucose and insulin tolerance testing**

Mice were fasted four hours starting in the morning (8–9am). Baseline (fasting) blood glucose was measured before insulin or glucose administration from venous blood via a small tail tip cut. Insulin (1U/kg body weight) (ITT) or Glucose (2 g/kg body weight) (GTT) (2 g/kg body weight) was injected intraperitoneally. Blood glucose values were obtained at 15, 30, 60, 90 and 120 min from the initial tail cut and measured using Contour® Next Blood Glucose Test Strips (Ascensia).

**Liver histology**

The liver was preserved in a 10% Neutral Buffered Formalin (VWR, 89370-094) for 72 hours at 4°C and then placed in a 70% ethanol solution. The samples were embedded in paraffin, cut, and stained by the Research Histology core at the Huntsman Cancer Institute (University of Utah) and ARUP Research Institute (Salt Lake City, Utah). Staining was for either Masson's trichrome or Hematoxylin and Eosin (H&E) staining.

**OCT liver preservation**

Liver tissue was either preserved directly in OCT solution (Tissue-Tek, 4583) or were fixed in 10% Neutral Buffered Formalin (VWR, 89370-094) for 24 hours before being dehydrated using a 30% sucrose solution for 48 hours at 4°C before being preserved in OCT solution. Preserved livers were frozen using dry ice and then stored at -80°C.

**Deparaffinization and rehydration**

Sections of human liver were fixed in 10% formalin (Sigma) and mounted into paraffin blocks. Tissue was then sectioned into 8 μm sections and mounted on glass cover slides. Sections were then deparaffinated by submersion in Histosol 2x for 10 min and rehydrated by submersion in a series of ethanol solutions for 5 mins (100%, 95%, 70%) followed by submersion in MilliQ water. Antigen retrieval was achieved by heating in Citrate buffer (40 mM Sodium Citrate Dihydrate, 60 mM Citric Acid, pH 6). Slides were submerged into boiling Citrate buffer for 10 minutes before being submerged in cold MilliQ water for 10 minutes.

**Immunohistochemistry**

Tissue sections were permeabilized in Tris-buffered saline (TBS: 150 mM NaCl, 50 mM Tris-HCl, pH 7.6) containing 0.2% Tween-20 and 1% BSA (Sigma, A6003) for 15 mins before being blocked with 10% Goat Serum (Invitrogen, 50062Z) containing 1% BSA for 2 hours at RT. Primary antibody (Novus anti-EFHD1, NBP2-33281) was diluted 1:200 in TBS containing 1% BSA. The diluted primary antibody was added to the slide and the slide was placed in a humidity chamber overnight at 4°C. The slides were then washed 2x in TBS and then incubated in TBS containing 0.3% $H_2O_2$ for 15 mins at RT. The secondary antibody (HRP-conjugated anti-rabbit, Jackson

Immuno Research, 111-035-144) was diluted 1:500 in TBS containing 1% BSA and added to the slides for 1 hour at RT in a humidity chamber. The slides were washed 3x in TBS and then treated with DAB Substrate (Thermo, 34002) for 5 mins. Slides were then washed 2x with TBS and 3x with MilliQ water before being stained with Mayer's Hematoxylin Solution for 1 minute. Slides were then washed again 3 x with 10 mM NaOH and 3x with MilliQ water before being covered with coverslips prior to imaging.

**Histology scoring**

Liver sections stained with either Masson's trichrome or H&E stain were scored by a liver pathologist (K.J.E.) who has blinded to the genotype and dietary condition. Livers sections were scored as follows(5, 6): Total steatosis (Total Fat), Macrovesicular steatosis (large droplet), Microvesicular steatosis (small droplet) and Hypertrophy (Ballooning): <5% [of field] = 0; 5-33% = 1; 34-66% = 2; >66% = 3. Inflammation: Number of inflammatory foci/field: <0.5 = 0, 0.5-1 = 1, 1-2 = 2, >2 = 3, where a focus is defined as a cluster of ≥ 5 inflammatory cells. MASH activity score was calculated for each mouse by adding the total steatosis, hypertrophy, and inflammation scores. Fibrosis was also scored as follows: No fibrosis = 0; Fibrous portal expansion = 1; Rare bridge of septae = 2; Numerous bidges or septae = 3.

**Lipid droplet size and count**

Liver H&E-stained sections were imaged and processed for noise reduction and background correction, then visualized with the same intensity ranges for comparison. Lipid droplet size and count were quantified as described previously using MATLAB R2023a (RRID:SCR_001622)(7).

**Nuclei clustering**

5-10 individual fields from each mouse H&E liver section were analyzed by an individual blinded to genotype and diet condition. Nuclei clusters, defined as any cluster of >3 proximal or touching nuclei, were counted manually.

**Fibrosis area**

5-10 individual fields from each mouse Masson's trichrome-stained liver section were imaged by an individual blinded to genotype and diet condition using an Olympus BX51WI microscope with a DP73 color camera and CellSens software at 10 X magnification. Colorized trichrome images were imported into ImageJ and the colors were isolated (pipeline: "Image">:"Color">"Split Channels"). This isolated the "blue", "green" and "red" color fields. We found that the "red" color field highlighted the fibrotic areas of the tissue most effectively. The "red' color fields were then exported into CellProfiler(8, 9) where the size of the fibrotic area was calculated using the IdentifyPrimaryObjects and MeasureObjectSizeShape modules. Settings were the same across all images. Measurements were exported to Excel for summary analyses.

**Immunofluorescence imaging**

OCT preserved liver tissue was sliced to 12 µm thick sections and placed on a glass microscope slide (VWR Superfrost Plus Slide, 48311-703). The sections were washed to remove OCT and then fixed in 10% Neutral Buffered Formalin for 15 minutes. Sections were then permeabilized in PBS containing 0.5% Triton-X 100 overnight in a humidity chamber with gentle agitation at RT. Sections were then washed in PBS and incubated with ImageIT FX Signal Enhancer (Thermo, I36933) for 30 minutes at RT. Sections were washed in PBS and blocked using goat serum for 90 minutes with gentle agitation at RT. Sections were then placed in primary antibodies (1:200 dilution in PBS containing 0.2 % Triton-X 100) and left overnight with gentle agitation at RT. Sections were washed in PBS then placed in secondary antibodies (1:500 dilution in PBS containing 0.2 % Triton-X 100) and left in the dark with gentle agitation at RT for 4 hours. Sections were washed in PBS and treated with DAPI solution (1:500 dilution in PBS containing 0.2 %

Triton-X 100, 300 uM stock from powder, Cayman Chemicals, 14285) for 2 minutes before being washed in MilliQ water. Sections were then left to dry in the dark for 5 minutes before being covered with a coverslip (Fisherbrand, 22x30-1, 12545A) using Prolong Gold Antifade solution (Thermo, P36980). Sections were left to cure in the dark overnight at RT before imaging.

#### Transmission electron microscopy

Mice were euthanized as described above and their livers dissected and minced into ~1 mm pieces. The liver was then fixed overnight at 4°C in Fixative solution (1% glutaraledehyde; 2.5% paraformaldehyde; 100 mM Cacodylate buffer, pH 7.4; 6 mM $CaCl_2$; 4.8% Sucrose). The next day, the liver was washed 3 times in Cacodylate buffer consisting of (mM) 200 sodium cacodylate salt, 2300 Sucrose, 16 $CaCl_2$) before undergoing secondary fixation in 2% Osmium tetroxide at room temp for 1 hour. Liver pieces were then washed twice in Cacodylate buffer and once in $ddH_2O$ before being stained with saturated uranyl acetate for 1 hour at RT. Liver pieces were then dehydrated with graded ethanol series (30%, 50%, 70%, 95%, 100% for 15 minutes each) and further dehydrated by incubating in absolute acetone 3 times for 10 minutes each. The liver pieces were then infiltrated with Epon epoxy resin using a graded epoxy series (30% for 5 hours; 70% for overnight; 100% for 8 hours with 3 changes; 100% fresh resin for embedding). The polymerization step was performed in fresh resin for 48 hours at 60°C. Ultracut was performed using Leica UC 6 ultratome, sections were taken at 70 nm thickness and mounted on 200 mesh copper grids. The grids with the sections were stained for 20 minutes with saturated uranyl acetate and 10 minutes with lead citrate. Sections were examined at an accelerating voltage of 120 kV in a JEOL-1400 plus (JEOL, Japan) transmission electron microscope equipped with CCD Gatan camera. Processing and imaging was performed by the University of Utah's Electron Microscopy Core. Distance and size measurements for mitochondrial size and ER-mitochondrial distances at contact sites were obtained using ImageJ by an individual blinded to treatment and genotype.

#### Hepatocyte isolation

Mice were fasted for 4 hours prior to procedure. Hepatocytes were isolated using a two-step isolation as follows: Hepatocyte wash buffer (HBSS [Thermo Fisher] supplemented with (g/L) 0.4 KCl, 1 D-glucose, 2.1 $NaHCO_3$, 0.2 EDTA), Calcium wash buffer (HBSS supplemented with (g/L) 0.4 KCl, 1 D-glucose, 2.1 $NaHCO_3$, 0.2 mM $CaCl_2$) and Liver Digestion Media (Hepatocyte Wash Media (Thermo, 17704024) supplemented with 1 mg/mL of Collagenase IV (Worthington CAT: LS004188)) were heated to approximately 42°C in a water bath. One end of a peristaltic pump was placed into the warm Hepatocyte wash buffer and the other end was affixed to a 25G needle (preferably blunted). Hepatocyte wash buffer was run through the tubing of the pump (ensuring that there were no air bubbles) and the flow was set to around 2-3 drops per second. The mouse was anesthetized using isoflurane and transferred to a perfusion tray with isoflurane hookup/nose piece and placed under a heat lamp. The mouse was confirmed as fully anaesthetized before proceeding by means of a toe pinch. The mouse was cut open from the groin to the chest, ensuring not to cut into the thoracic cavity. The abdominal fat and intestine were gently moved aside using blunt forceps until the inferior vena cava (IVC) was exposed. The 25G needle was then gently threaded into the IVC ensuring that the tip of the needle was above the renal branches but did not protrude into the liver and clamped. The peristaltic pump was switched on and the liver was observed until distinct lighter spots began to appear on the lobes, indicating that the lobes were filling with Hepatocyte wash buffer. At this point, the portal vein was cut to allow blood and solution to exit the liver. The solution passing through the pump was switched to Liver Digestion Media, ensuring that no bubbles entered the tubing and was allowed to run for 10 minutes. In our hands, this time allowed for ~15 mL of Liver Digestion Media to pass through the liver. After 10 minutes, the needle and clamp were removed and the liver was gently dissected from the surrounding tissue and placed in a 10 cm dish containing Liver Digestion Media. The liver was gently broken apart using blunt forceps until it formed a soup-like consistency before being transferred together with the Liver Digestion Media to a conical flask and agitated at 37°C for 60 mins. The liver remnant was then filtered through a 100 μM nylon mesh filter and centrifuge at 50 *x g* for 5min at RT (for MASH-fed animals, this was increased to 150 *x g*). The supernatant was removed and the pellet was suspended in Calcium wash buffer and spun as above before being suspended again in Modified Williams Media E (Williams Media E (Thermo, 12551032) supplemented with 1% (v/v) of the following: glucose solution (A2494001), NEAA (1140050), Sodium Pyruvate (11360070), HEPES solution (15630080), Glutamax (35050061), Pen/Strep (15140148) and 10% (v/v) FBS (Thermo, 26-140-079)). Cells were then plated at a low density (~30% confluence, to preserve viability) on gelatin-coated coverslips or cell culture plates and left overnight at 37°C with 5% $CO_2$.

**Live cell imaging of $Ca^{2+}$ uptake and ATP- and ionomycin-induced fission**

Hepatocytes, HepG2 or HAP-1 cells were treated with 200 nM MitoTracker Orange (Thermo, M7510) for 30 minutes at 37°C. For direct measurement of fission events in HepG2 cells, these were transfected with mitochondrially-targeted mGold2s (pCMV-mGold2-Mito-7, gift from the St. Pierre Lab)(10) for time-course HepG2 cell imaging. For live actin imaging, cells were stained with SiR-Actin (VWR, CY-SC001) for 30 minutes at 37°C. For obtaining $Ca^{2+}$ transient data, cells are stained with 1 μM Cal520-AM (AAT Bioquest, 21130) which senses cytosolic $Ca^{2+}$ and 5 μM X-rhod-1 AM (Therom, X14210) which senses mitochondrial $Ca^{2+}$ for 30 minutes at 37°C. Imaging took place in the cell media using a confocal microscope with a heated chamber at 37°C with 5% $CO_2$ (Nikon Eclipse Ti2 microscope with Photometrics Prime BSI scientific CMOS camera and okolab incubator) or at room temperature (Leica Microsystems SP8 microscope, Germany). For ATP experiments, freshly made 100 mM ATP stock was prepared in PBS and was added to the plate media and allowed to incubate for 5 minutes before imaging. For ionomycin experiments, 1 μM ionomycin in DMSO was added to the plate and allowed to incubate for 5 minutes before imaging.

**Mitochondrial length measurement**

Images of cells stained with MitoTracker were processed in ImageJ(8) to remove noise and enhance mitochondrial staining using a bandpass filter. Images were then exported to either CellProfiler(9) or iLastik (version 1.4.1; https://www.ilastik.org)(11). The ImageJ, CellProfiler, and iLastik pipelines were applied identically to all images regardless of genotype or condition. For analysis of mitochondria in HepG2 or HAP-1 cells we used CellProfiler. The pipeline sequentially applied the GaussianFilter, EnhanceOrSuppressFeatures, IdentifyPrimaryObjects, SplitOrMergeObjects, MeasureObjectSizeShape, and export modules.

Mitochondrial staining in isolated hepatocytes was too dense for accurate measurement with CellProfiler alone, so these were segmented and quantified using the interactive machine learning software iLastik. We employed the Pixel Classification workflow to segment mitochondria. Training was performed locally on a workstation by manually annotating representative regions of interest. Using ilastik's brush tool, we labeled pixels corresponding to either mitochondria or background across a subset of images. Feature selection included intensity, edge, and texture-based descriptors at multiple spatial scales. The classifier was iteratively refined using ilastik's "Live Update" mode, which provides real-time feedback on segmentation quality. To assess segmentation accuracy, we visually inspected overlay images of the raw data and predicted masks. After training the classifier, all images across conditions and genotypes were segmented identically, and mitochondrial area and length were measured using ilastik's in-built Object Classification workflow. This step segments individual mitochondria as discrete objects and allowed for the measurement of properties like area, shape, and intensity. The measurements were then exported to an Excel spreadsheet for analysis.

**Counting fission events in live cells**

Isolated mouse hepatocytes treated with MitoTracker or HepG2 cells transfected with mito-mGold2s were imaged as described above (Live cell imaging). Imaging took place over three minutes with images taken every 1 second. A fission event was defined as the separation of a larger mitochondrial structure into two smaller structures as described previously(12) . Fission events were counted manually by an individual who was blinded to genotype and treatment.

#### $Ca^{2+}$ induced fission events in permeabilized mitochondria

Hepatocytes were isolated as described above and incubated with 200 nM MitoTracker orange. Media was removed and replaced with imaging solution consisting of (mM) 125 KCL, 20 HEPES, 5 $K_2HPO_4$ , 1 $MgCl_2$ (pH to 7.2 with KOH, osmolality 290–300 mOsm/L)) supplemented with 1 μM Ru360 and 60 μM EGTA. Cells were imaged every minute for 6 minutes. At t=1 minute, 0.002% (w/v) digitonin was added to the cells to permeabilize the cells. At t=2 minutes, 0.1 mM $CaCl_2$ was added to the cells. Mitochondrial length was calculated at each minute as described above.

#### Hepatocyte starvation

Hepatocytes were isolated as described above and incubated for 24 hours in either “starvation” media (1 g/L D-glucose DMEM, Gibco 11885-084, Life Technologies) or “fed” media (4.5 g/L D-glucose DMEM, Gibco 11965-092, Life Technologies, supplemented with 0.4 mM palmitate conjugated to BSA) before being imaged live as described above.

#### Mitochondria isolation

Mice were euthanized as described above and the organs were removed and washed in ice-cold phosphate buffered saline (PBS). Livers were rapidly dissected and placed in ice-cold initial medium containing (mM): 225 mannitol, 70 sucrose, 5 HEPES, 1 EGTA, and 0.1% (w/v) bovine serum albumin (BSA) (pH to 7.2 with KOH; osmolality to 290–310 mOsm/L). Tissue was homogenized using a Potter-Elvehjem tissue grinder attached to an overhead stirrer (IKA, Wilmington, NC) for 10–15 strokes at 180 rpm. The homogenate was centrifuged at 700 × *g* for 7 minutes and the tissue pellet was discarded. The supernatant was then centrifuged at 9000 × *g* for 9 minutes to obtain a mitochondrial fraction. The mitochondrial pellet was washed twice with and carefully resuspended, using a pipette tip where the end had been removed, in imaging solution containing (mM): 125 KCL, 20 HEPES, 5 $K_2HPO_4$, 1 $MgCl_2$, and 0.01 EGTA (pH to 7.2 with KOH, osmolality 290–300 mOsm/L). Protein concentration was quantified using the Pierce 660 nm Protein Assay Reagent (Thermo Fisher, Waltham, MA) according to the manufacturer’s directions.

#### Isolation of mitochondrial-ER contact sites (ERMCSs)

ERMCSs were isolated as described previously(13). Briefly, the crude mitochondrial pellet was obtained as described above (Mitochondria isolation). The crude mitochondria were then layered on top of Percoll medium, consisting of (mM): 225 mannitol, 25 HEPES, 1 EGTA, 30% Percoll (pH 7.4). The crude mitochondria and Percoll medium were spun at 95,000 × *g* for 30 minutes at 4°C. A dense band containing pure mitochondria was located at the tube bottom, while a dense band containing ERMCSs was visible above this mitochondrial band. These bands were extracted and separately spun at 6,300 × *g* for 10 minutes at 4°C. The mitochondrial fraction was then collected. The ERMCS fraction was spun again at 100,000 × *g* for 1 hour using a Beckman Coulter Opima MAX-XP Ultracentrifuge with an MLA-55 rotor holding 16 X 76 mm Ultra-Clear open-top centrifuge tubes (Beckman Coulter), and the pure ERMCSs collected. Purity of each fraction was confirmed via Western Blot using antibodies against known organelle marker proteins.

#### Western blot

Tissue was lysed in RIPA lysis buffer supplemented with protease and phosphatase inhibitors (Thermo Fisher) and then mixed in a 1:1 ratio with Western blot loading dye (4X Bolt LDS Sample

Buffer supplemented with 10 mM Dithiothreitol (DTT). Protein concentration was quantified by BCA assay (Thermo Fisher). 5–20 µg of protein from total heart lysates were loaded on polyacrylamide gels and processed as described previously on PVDF membranes(14). Antibodies used are listed above. Band intensity was analyzed using ImageJ(8). Expanded views of all membranes are available in Figure S10.

**Co-immunoprecipitation**

HepG2 cells were grown and transfected with either an EFHD1-HA construct, or an OMP25-GFP-HA construct (a gift from David Sabatini, Addgene plasmid # 83356)(15) . Cells were then harvested and lysed in a PBS solution containing 1 % Triton-x-100 and 10 mM phenylmethylsulfonyl fluoride (PMSF) by passing through a 25G needle. The lysates were quantified using a BCA assay and diluted to 250 µg/mL of protein. 500 µL of the lysates were agitated in 20 µL of EZview™ Rd Anti-HA Affinity Gel beads (Sigma, E6779) overnight at 4°C. The beads were washed four times in PBS+1% Triton-X-100 before being mixed 1:1 with western blot loading dye and loaded on a gel for western blot analysis.

**AlphaFold3 modeling**

EFHD1-actin structures were predicted computationally, as we have done previously, using the publicly-available AlphaFold3 server(16, 17). Sequences used in the manuscript were for 1 human EFHD1 (Uniprot Q9BUP0), 1 human β-actin (Uniprot P60709), 2 $Ca^{2+}$ ions, and 1 ATP molecule. Interface predicted template modeling (ipTM) and predicted local distance difference test (pLDDT) metrics were used to judge the quality of the prediction. The top 5 predicted structures were visualized on ChimeraX(18). Predicted actin-EFHD1 structures were compared to actin-fimbrin (3BYH)(19), actin-fascin (8VO6)(20), and actin-filamin (6D8C)(21) cryo-EM structures. Structures with other stoichiometries or without $Ca^{2+}$ or ATP all had actin and EFHD1 in the same interaction pose, but with slightly lower ipTM values (some below the 0.8 cutoff), so we did not analyze these further.

**Whole mitoplast electrophysiology**

Whole-mitoplast electrophysiology was performed as described previously(22-24). Whole mitoplast currents were measured using patch clamp. Giga-ohm seals with mitoplasts were formed in the KCl bath solution. Voltage steps of 350–500 mV for 2–8 ms were applied to rupture the IMM and obtain the whole-mitoplast configuration. Typically, pipettes with 25–35 MΩ resistance were used for the patching. Currents were normally induced by a voltage ramp from −160 mV to +80 mV (interval between pulses was 5 s). All whole-IMM recordings were performed under continuous perfusion of the bath solution. Pipettes were filled with internal solution containing (in mM): 110 Na-gluconate, 40 HEPES, 10 EGTA and 2 MgCl2 (pH 7.0 with NaOH) (tonicity was adjusted to ~350 mmol/kg with sucrose). To measure whole-mitoplast $Ca^{2+}$ currents, the bath solution contains (in mM): 150 HEPES, 80 sucrose, and 1 $CaCl_2$ (pH 7.0 with Trizma base, tonicity ~300 mmol/kg with sucrose). Currents were normalized per membrane capacitance to obtain current densities (pA/pF). For display purposes, capacitance transients caused by changing levels of solutions in the bath have been removed. Analysis was performed using pClamp v10 (Molecular Devices).

**RNA-seq methods and analysis**

Normal chow and MASH diet RNA-seq experiments were done separately. Mice were euthanized as described above and the liver dissected, homogenized and stored in TRIzol™ Reagent (Thermo, 15596026) RNA was isolated using the Direct-zol™ RNA MicroPrep kit (Zymo, R2062) after which RNA was submitted to the Huntsman Cancer Institute High-Throughput Genomics Shared Resource center for library preparation and Illumina sequencing. Total RNA samples (100-500 ng) were hybridized with Ribo-Zero Gold to substantially deplete cytoplasmic and mitochondrial rRNA from the samples. Stranded RNA sequencing libraries were prepared as described using the Illumina TruSeq Stranded Total RNA Library Prep Gold kit (20020598) with TruSeq RNA UD Indexes (20022371). Purified libraries were qualified on an Agilent Technologies 4150 TapeStation using a D1000 ScreenTape assay (cat# 5067-5582 and 5067-5583). The molarity of adapter-modified molecules was defined by quantitative PCR using the Kapa Biosystems Kapa Library Quant Kit (cat#KK4824). Individual libraries were normalized to 0.65 nM (normal chow) or 1.30 nM (MASH diet). Sequencing libraries were chemically denatured and applied to an Illumina NovaSeq flow cell using the NovaSeq XP workflow (20043131). Following transfer of the flowcell to an Illumina NovaSeq 6000 instrument, a 151 x 151 cycle paired end sequence run was performed using a NovaSeq 6000 S4 reagent Kit v1.5 (20028312). The pipeline protocol was single-base mismatch.

For normal chow, the mouse GRCm38 (mm10) genome and gene annotation files were downloaded from Ensembl release 102 and a reference database was created using STAR version 2.7.6a(25). For MASH diet, the mouse GRCm39 genome and gene annotation files were downloaded from Ensembl release 110 and a reference database was created using STAR version 2.7.9a(25). Optical duplicates were removed from the paired end FASTQ files using clumpify v38.34 and reads were trimmed of adapters using cutadapt 1.16(26). The trimmed reads were aligned to the reference database using STAR in two pass mode to output a BAM file sorted by coordinates. Mapped reads were assigned to annotated genes using featureCounts version 1.6.3(27). The output files from cutadapt, FastQC, FastQ Screen, Picard CollectRnaSeqMetrics, STAR and featureCounts were summarized using MultiQC to check for any sample outliers(28). Differentially expressed genes were identified using a 5% false discovery rate with DESeq2 version 1.30.0 (normal chow) or 1.40.2 (MASH diet)(29). Significantly enriched Hallmark, KEGG and REACTOME pathways were detected using the fast gene set enrichment package(30). The sequencing data has been deposited in the NCBI Gene Expression Omnibus database.

**Mass Spectrometry Sample Preparation**

Mice were euthanized as described above and the liver dissected, homogenized and lysed in RIPA buffer. Proteins from the samples were digested as previously published(22). 10 µg from each sample was loaded onto separate Vivacon 500 filter units, concentrated at 13,000 x *g*, and then washed three times with 100 µL of urea buffer (8M Urea, 0.1M Tris/HCl pH 8.5). The concentrate was then mixed with 100 µL of 50 mM iodoacetamide in urea buffer and incubated at room temperature in the dark for 20 minutes, followed by centrifugation for 15 min at 13,000 x *g*. The concentrate was then washed twice with 100 µL of urea buffer and two washes with 100 µL of 50 mM ammonium bicarbonate. 10 µg of protein from each sample was mixed with trypsin (1:40) in the filter and incubated overnight at 37°C. The peptides were eluted with 50 mM ammonium bicarbonate and acidified with 1% formic acid.

**Mass Spectrometry Acquisition and Analysis**

Three technical replicates were used for each mouse sample (biological replicate). Tryptic peptide samples were analyzed on a Vanquish Neo UHPLC system (Thermo Fisher Scientific) coupled via a nano-electrospray ion source to an Orbitrap Astral mass spectrometer (Thermo Fisher Scientific). About 500 ng of peptides were loaded onto an IonOpticks Aurora TS C18 analytical column (75 µm × 25 cm, 1.7 µm, 100 Å) using buffer A (0.1% formic acid in water). Separation was achieved with a linear gradient from 4% to 22.5% buffer B (80% acetonitrile, 0.1% formic acid in water) over 12 min, followed by an increase to 40% buffer B over 7.5 min, and a final ramp to 99% buffer B over 5.5 min. Data were acquired in data-independent acquisition (DIA) modes. For DIA, full MS scans were acquired in the Orbitrap at 240k resolution with 500% automated gain control (AGC) and a precursor mass range of 380–980 m/z. DIA MS/MS scans were performed in the Astral at 80k resolution with a fixed 2 m/z isolation window, a maximum injection time of 3.5 ms, and higher-energy collisional dissociation (HCD) at 25%, with AGC at 500% over a scan range of 150–2000 m/z. All MS data were searched against the UniProt mouse reference proteome (17,090 protein entries) to identify matching tryptic peptides. DIA data were processed in Spectronaut (version 19.5, Biognosys). Searches allowed up to two missed tryptic cleavages, with mass tolerances set to 20 ppm for precursor ions and 0.02 Da for fragment ions. Cysteine carbamidomethylation was specified as a fixed modification, while N-terminal acetylation and methionine oxidation were set as variable modifications. Precursor ion charge states were restricted to +2, +3, and +4. All searches were performed against a target-decoy database, and protein identifications were filtered at a 1% false discovery rate (FDR). All data was analyzed using Perseus (version: 2.0.11.0) and R studio (version: 2024.04.1). The mass spectrometry raw files have been uploaded to the PRIDE database via the PRIDE partner repository. The proteomics data were then analyzed for REACTOME pathway enrichment with the PADOG pipeline or fast gene set enrichement(30-32). Selected pathways from either analysis were chosen for display using the average fold change calculated from PADOG, with the false discovery rate coming from the analysis from which the pathway was chosen. Differential expression analysis was via the limma package(33). For correlation plots, average fold changes calculated from PADOG analysis for the KO MASH versus WT MASH comparison were plotted against the average fold change of the WT GAN versus WT CHOW comparison. Correlations were for all pathways across the entire REACTOME map, or for those pathways contained within the Immune System (R-HSA-168256), Fatty acid metabolism (R-HSA-8978868), or Translation (R-HSA-72766) nodes.

**Isolation of double-stranded RNA and PKR-bound RNA**

Immunoprecipitation via dsRNA and PKR antibodies was modified from a protocol described previously(34). Briefly, hepatocytes were isolated as described above and fixed in 0.1 % formalin in modified Williams Media E at room temperature for 10 mins before being quenched by adding 125 mM glycine in PBS. Hepatocytes were spun down and sonicated for 2s on and 2s off for 5 mins in in 600 µL fCLIP lysis buffer consisting of (mM) 20 Tris-HCL [pHn 7.5], 15 NaCl, 10 EDTA, 0.5% NP-40, 0.1% Triton X-100, 0.1% Na-deoxycholate). Hepatocytes were then incubated in either dsRNA antibody (15 µL, clone rJ2, Sigma, MABE1134), PKR antibody (50 µL, Santa Cruz, sc-6282) or mouse IgG (15 µL, Normal Mouse IgG, Sigma, 12-371) for 1 hour before 20 µL of magnetic beads (Dynabeads™ Protein G for Immunoprecipitation, Thermo, 10003D) were added to each tube and incubated for a further 2 hours. Hepatocytes were washed 3x with fCLIP (1 mL) and 1x with EB (100 µL) before being eluted with Urea buffer consisting of (mM) 4000 Urea, 200 Tris-HCL [pH 7.4], 100 NaCl, 20 EDTA, 2% SDS). The elutes were mixed with 100 µL of proteinase K (20 mg/mL) and left overnight at 65C. The next day the eluates were mixed with 1 mL TRIzol and 200 µL of chloroform was added to each sample and incubated on ice for 5 mins. Samples were spun at 4C at 13 000 x g for 5 mins and the upper aqueous phase (~480µL) as above into equal volume of isopropanol. 2 µL of GlycoBlue (thermo) was added to each sample and inverted by hand 10 times to mix and incubated at -20°C for 20 min. The samples were spun at 4°C at 13 000 x g for 5 mins to precipitate RNA and the supernatant was discarded. The RNA precipitate was washed with 70% ethanol and left to air-dry at for 5 mins. The RNA was resuspended in 15-20 µL of RNAse-free water and then mixed with 500 µL TRIzol for storage (Thermo, 15596026).

**Quantitative reverse transcriptase polymerase chain reaction (qRT-PCR)**

RNA samples stored in TRIzol were further purified using the Direct-zol™ RNA MicroPrep kit (Zymo, R2062). RNA was converted to cDNA using SuperScript™ IV VILO™ Master Mix (Thermo, 11756050). 15 ng of cDNA was mixed with 300 nM of primer pairs (designed using

Primer-BALST) in qPCR mix (SYBR™ Green PCR Master Mix, Thermo,4309115) analyzed via qPCR analysis using a CFX Opus 96 (Biorad)

Data was exported to Excel and fold change was calculated by the -2ΔΔCt method.

**UK Biobank (UKB) study cohorts and data**

The UKB is a population-based prospective study established to allow detailed investigation of the determinants of disease in middle to old age adults(35). It includes approximately 500,000 participants, aged 40 to 69 years recruited between 2006 and 2010, with comprehensive clinical and laboratory data as well as genome-wide genotyping data(36). All participants of the UKB provided informed consent. We focused our analyses on 461,621 unrelated individuals with available serum levels of AST from their initial visit and measured centrally by the UKB.

**Genetic data, imputation, and quality control**

Genome-wide genotype data for UKB participants was accessed through the UKB Research Analysis Platform. Briefly, genotyping of approximately 826,000 variants was performed using either Affymetrix's UK BiLEVE Axiom array or Applied Biosystems' UK Biobank Axiom array. Imputation to >93.1 million variants was performed using the TOPMed reference panel and has been described previously(36). Variant QC procedures, including filters for minor allele frequency (MAF) ≥ 0.01, Hardy-Weinberg equilibrium p-value <1e-15, individual missingness ≥10%, genotype missingness ≥10%, and linkage disequilibrium (LD) pruning, were implemented, yielding 5,024,228 high-quality variants. Imputed data underwent additional QC based on imputation quality (INFO score ≥0.4). To ensure independence in genetic analyses, related individuals were identified using KING software(37) and individuals who were first-degree relatives (kinship coefficient >0.125) were excluded. Genetic ancestry of UKB participants was estimated using principal components analysis as implemented in PLINK2(38).

**Association of variants at *EIF2AK2* (*PKR*) with AST levels and liver eQTLs**

Association analyses of variants at the *EIF2AK2* locus (spanning a 2,057,771 basepair region from positions 36089210 to 38166980 on chromosome 2 (based on human genome build hg38), which included a total of 5,635 variants located 1 mega-basepair up- and down-stream of the *EIF2AK2* reference sequence (NM_002759.4)). Association analyses were performed using REGENIE version 3.3, which implements a two-step machine learning approach designed to handle large-scale genomic data, on rank-based inverse normal transformed AST levels and included adjustment for age, age squared, sex, BMI, and the first 10 principal components of genetic ancestry(39). Variants achieving a Bonferroni corrected p-value < $8.87 \times 10^{-6}$ (0.05/5,635) were considered statistically significant.

Colocalization analyses were then conducted to investigate the shared genetic architecture between variants at the *EIF2AK2* locus and liver expression quantitative trait loci (eQTL) and to identify genetic signals that may influence *EIF2AK2* gene expression using liver eQTL meta-analysis data from 1,183 individuals(40). Additionally, variants with liver eQTLs were annotated using RegulomeDB, version 2.2(41). In brief, RegulomeDB assigns a ranking score to each variant, based on the variant's location in one or more of the following: eQTL, transcription factor (TF) binding, TF motif, DNAse footprint, and DNAse peak, denoting its likelihood of being functional, and also calculates a model prediction score to predict the probability of the query variant being functional using a random forest model.

**Association of variants at *EIF2AK2* (*PKR*) with liver disease diagnoses**

For variants significant from the above analyses, we performed association analyses for 406,863 participants with three tiers of liver disease diagnoses listed in UKB. Association analyses with were performed using REGENIE version 3.3, which implements a two-step machine learning approach designed to handle large-scale genomic data, on binary disease diagnosis trait and included adjustment for age, age squared, sex, BMI, and the first 10 principal components of genetic ancestry.(39) Three separate analyses were completed depending on liver disease inclusion: a broad category for all liver diagnoses in levels 1-3 (Tier 1, cases = 12,635, **Table S9**), a category excluding any cases with diagnoses associated with infectious causes, (Tier 2, cases = 11,839, **Table S10**), or a category for cases with chronic liver disease codes excluding infectious causes (Tier 3, cases = 4,635, **Table S11**).

**Mendelian Randomization (MR)**

One-sample MR was completed with UKB GWA summary statistics using the R package MendelianRandomization (version 0.10.0)(42) to investigate the causal role of variants associated with AST at the *EIF2AK2* locus on liver disease, using pQTL-associated variants as genetic instrumental variables (IVs). In MR analyses, genetic variants classified as IVs must satisfy three assumptions: 1) the genotype is associated with the exposure; 2) the genotype is associated with the outcome only through the exposure; and 3) the IV is independent of other factors that affect the outcome(43). Exposure data was obtained from GWA summary statistics for pQTL-associated variants. Three separate analyses were conducted with outcome data varying by liver diagnosis inclusion (**Table S7**). Exposure and outcome data were harmonized yielding 35 SNPs to be included in the MR analyses.

We prioritized exposures as causal when there was a significant effect in using a Bonferroni-adjusted p-value threshold $<1.42 \times 10^{-3}$ (p-value <0.05/35) for the inverse variance weighting.

**Statistics**

Microsoft Excel, OriginPro (OriginLab) and R Studio (Posit Software) were used for data analysis. For two-sample comparisons where n < 15 or where a Shapiro-Wilk normality test found a non-normal distribution, we used a Mann-Whitney non-parametric test. For comparisons on samples with normally distributed data sets, we used a two-tailed, unequal variance, Student's t-test. For comparisons of multiple-samples, a 1-way ANOVA with a Bonferroni-corrected post-test was performed for normal data and a Kruskal-Wallis ANOVA with a Dunn's multiple comparison test for non-normal data. For analysis of fluorescence data sets consisting of repeated measures from individual mice (e.g. mitochondrial lengths) we utilized a Linear Mixed Effects Model in R to account for the hierarchical nature of the data, as described(44).
